# Insights into the functional and genetic basis of heteranthery in *Arthrostemma ciliatum* Pav. ex D.Don (Melastomataceae)

**DOI:** 10.64898/2026.02.02.703258

**Authors:** Suvrat Kotagal, Anna Schlick, Christian Siadjeu, Emy Yue Hu, Gudrun Kadereit

## Abstract

**Background:** Melastomes are well known for their striking diversity in stamen morphologies mostly adapted to buzz pollination by bees. The variously modified connective appendages and heteranthery in the family have fascinated botanists for more than two centuries and a variety of functions associated with pollination have been discovered for these staminal traits over the years. The repeated evolutionary shifts in these traits have been linked to pollinator shifts, likely contributing to diversification in the family. The evolutionary lability of staminal traits, especially the connective morphology, led us to hypothesize that these traits might be controlled by relatively simple genetic mechanisms and we here take the first steps to test this hypothesis by using a comparative transcriptomics approach with *Arthrostemma ciliatum* as our model. We also tested the functional significance of heteranthery and whether the classical division of labour hypothesis holds true for this species by comparing the number, size and viability of pollen in the two stamen types.

**Results:** Staminal development of this species was studied and suitable stages for transcriptome comparisons were identified. Differential expression analyses between the morphologically distinct stamen whorls at four developmental stages showed the differential expression of several transcripts involved in stamen development/elongation. Pollen comparisons between the two whorls showed that the antepetalous/inner whorl stamens have a significantly higher number of pollen and higher germination rates while the antesepalous/outer whorl stamens have significantly larger pollen.

**Conclusions:** We identified Jasmonate and Gibberellin signalling pathway genes (*JAZ*, *GID1*, *DELLA* and *ARF* homologs), *EPF*/*EPFL* family genes, autophagy related genes (*VPE* homologs) and S Locus ELF homologs as putative candidates involved in causing staminal dimorphism in *A. ciliatum.* Our results indicate that, for the heterantherous morph of this species, the shorter stamens (antepetalous/inner whorl) have both pollinating and feeding functions contradicting the division of labour theory. We also report the possible existence of heterostyly in *A. ciliatum* as an outbreeding mechanism.

## Introduction/Background

Heteranthery (also called staminal dimorphism) is the presence of stamens of different size, shape and/or colour in a flower. This phenomenon has evolved several times independently and is known in at least 20 angiosperm families [1] prominent among them is the pantropically distributed and species-rich Melastomataceae [2] which show a striking diversity in androecial traits [3]. Heterantherous flowers are often nectarless and associated with animal pollination; especially buzz pollination with bees being the effective pollinators [1]. A plausible explanation for the function of heteranthery suggested by H. & F. Müller [4, 5] is the “division of labour hypothesis”. It states that the two different kinds of stamen perform different functions in the flower – one set provides the pollen reward and attracts pollinators while the other set secures cross-pollination. However, some studies challenge this view [6] and provide alternative explanations, and the biological reality seems much more complex in light of the diversity of heterantherous phenotypes.

In Melastomataceae the morphological diversity of the androecium and the disparity in stamen types has been repeatedly hypothesised to be a factor involved in pollinator shifts and diversification in the family [7–13]. The astonishing heterogeneity in connective appendages and the constant shifts between iso-dimorphism of stamens in closely related taxa has been a focus of discussion in this family for many years. In particular, the ‘pedoconnective’ (*sensu* Jacques-Félix [14]- a basally elongated connective) has been considered a key innovation responsible for high levels of species diversity in certain clades of the family [10]. Many previous classifications of Melastomataceae have assigned taxonomic value to these staminal variations, albeit at different levels [9, 15–19].

Usually heteranthery in melastomes occurs between the two stamen whorls within a flower. Most commonly the stamens of the outer whorl (opposite to the sepals - antesepalous stamen) are significantly longer than the stamens of the inner whorl (opposite to the petals - antepetalous stamen). The morphological differences between the stamen whorls are gained during the course of development underlain by differences in organ initiation patterns and space constraints [20]. But heteranthery can also sometimes be seen within the same stamen whorl while the other is completely reduced as in *Rhynchanthera grandiflora* [21]. The developmental aspects of heteranthery are discussed in detail by Basso-Alves et al. [20] who emphasize that the diversity in stamen appendages is a consequence of distinct ontogenetic processes. It is clear that the phenomenon of heteranthery is - at least in a developmental sense - not homologous among melastomes and the existence of many intermediate phenotypes indicates that the underlying ontogenetic processes are labile.

Heteranthery in melastomes is generally attributed to the above mentioned division of labour hypothesis, where one set of stamens function as feeding stamen and the other set are involved in effective pollination in order to alleviate the ‘pollen dilemma’ (reduced male fitness due to pollen consumption) in buzz pollinated flowers [22–25]. The feeding stamen are often yellow to attract bees, while the longer pollinating stamen are camouflaged against the petals [22, 26, 27]. The pollinating stamen are positioned in such a way that they can maximise the area of pollen deposition [21] or place pollen on “safe sites” (out of reach for grooming) on the bees’ body [22, 28]. Connective appendages of the pollinating stamens can also mimic the feeding stamen, increasing the attractive potential by deceiving the pollinator [22, 27–29]. Additionally, there may exist differences in pollen number [27, 28] and morphology [22, 23] between the two stamen types (see also [30]). Some taxa invest more in feeding pollen to ensure pollinator visits [29] while others economize and invest more in pollinating stamen [27, 28]. Heteranthery has also been shown in non-pollen rewarding flowers, where gradual maturation of food bodies staggers pollen release and promotes repeated pollinator visits, enhancing outcrossing opportunities [31].

Considering how pedoconnectives and heteranthery are used by melastomes to successfully achieve pollination, it is not surprising that the variation in these characters is often associated with diversification in the family. Staminal traits including connective morphology, dimetrism and dimorphism show a high degree of evolutionary lability in the family [7, 10, 20, 32], especially among members of the so called ‘pedoconnective clade’ [3, 20]. The instability of these staminal traits between closely related species leads us to hypothesize that these traits are governed by a small set of genes that are subject to constant evolutionary tinkering. If so, this could not only explain the diversity of staminal traits in Melastomataceae but also represent a major driver of diversification of the family as even small changes in stamen morphology influence pollination systems [33].

The lability of heteranthery among melastomes is well exemplified by the small genus *Arthrostemma* Pav. ex D.Don within the pedoconnective clade. It belongs to the tribe Rhexieae, consisting of only four currently accepted species native to Mexico, Central America and Northern parts of South America [34]. Out of these, two have iso/subisomorphic stamen (*A. alatum* Triana and *A. parvifolium* Cogn.), one has dimorphic stamen (*A. primaevum* Almeda), while the fourth species (*A. ciliatum* Pav. Ex D.Don) is, to our knowledge, one of only two melastome species known to exhibit multiple staminal forms and the only species with both isomorphic and dimorphic flowers (Figure 1) [34–36]. Due to the co-occurrence of these two floral morphs in close geographic proximity (Figure 1), *A. ciliatum* is a particularly suitable system to study the evolutionary shifts of staminal traits.

**Figure 1:**
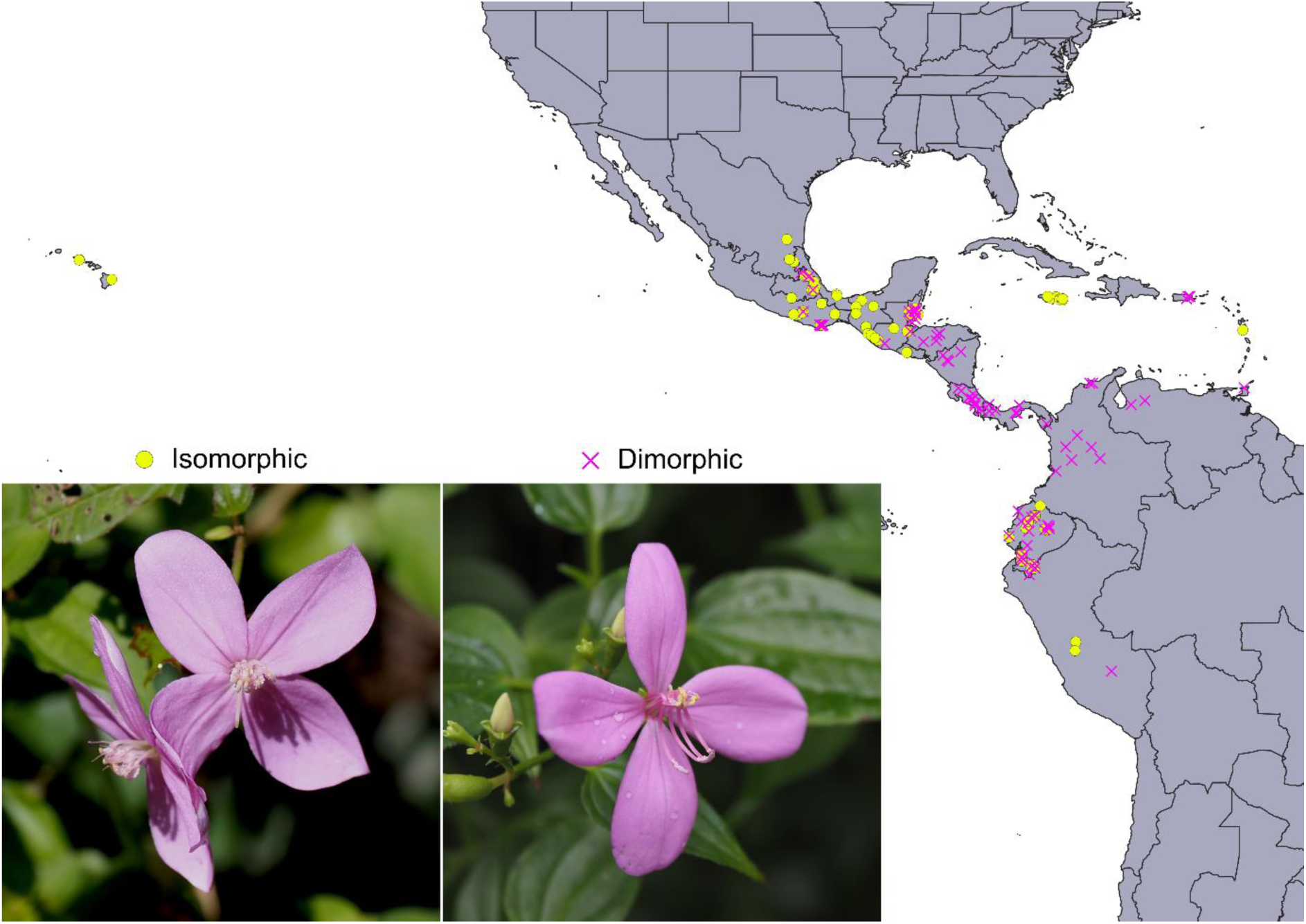
Distribution of the two stamen morphs of Arthrostemma ciliatum. Occurrence points are from specimens on the NYBG virtual herbarium website (https://sweetgum.nybg.org/science/vh/) and from inaturalist observations; only those coordinates are shown where the floral morph was clearly distinguishable from the images of the specimen/observations. Pink crosses on map refer to dimorphic individuals while yellow dots refer to isomorphic individuals. Inset shows flowers of the two stamen morphs. Isomorphic: Two flowers from the same inflorescence with different style positions and short stamen only. The style declinates later during anthesis (Image: S. Kotagal Pichincha, Ecuador). Dimorphic: Flower with long and short stamen. Style always close to the short stamen (Image: S. Kotagal, Botanical Garden Munich Nymphenburg, Germany).

Despite its ecological and evolutionary significance, the genetic basis of heteranthery is largely unknown. To date there are only three studies that address this knowledge gap and provide only the first insights into the topic. Studies on *Cassia biscapsularis* (Fabaceae) [37], *Melastoma dodecandrum* (Melastomataceae) [38] and *Monochoria elata* (Pontederiaceae) [39] each identified different candidate gene groups responsible for heteranthery with little overlap. This suggests that distantly related species have distinct genetic bases for heteranthery which again points to the multiple evolutionary origins of this convergent trait and highlights the need for further investigation, especially in the species rich Melastomataceae due to reasons outlined in the previous paragraphs

In this study we are setting a foundation for further studies on the evolution and genetic basis of heteranthery in melastomes by focusing on the dimorphic form of *A. ciliatum*. We aim to 1) characterize stamen development and select suitable stages for transcriptomic profiling, 2) identify candidate genes responsible for staminal dimorphism in *A. ciliatum* via differential expression analyses, and 3) test the division of labour hypothesis for *A. ciliatum* by assessing and using the number, size and viability of pollen in the two stamen types as a proxy for male fertility. We used developmental, genetic and functional analyses to provide insights into the mechanisms underlying heteranthery and its functional significance in *A. ciliatum*.

## Methods

### Developmental studies

The stamen development of the dimorphic *Arthrostemma ciliatum* was studied in order to identify suitable stages for transcriptome comparisons. Fresh (unfixed) flower buds from plants at the Botanical Garden Munich Nymphenburg (garden accession no. 2005/2472-1, herbarium specimen deposited at M) were collected and dissected across developmental stages, from the smallest hand-dissectible buds (∼0.5mm) to open flowers, and examined using a Keyence VHX-7000 digital microscope. The sizes of the flower buds at each developmental stage and the differences between the two stamen whorls (outer - antesepalous stamen, inner - antepetalous stamen; used interchangeably throughout this study) were documented. Four developmental stages were chosen based on their sizes and consistent morphological differences to perform transcriptomic comparisons (Table 1).

**Table 1:**
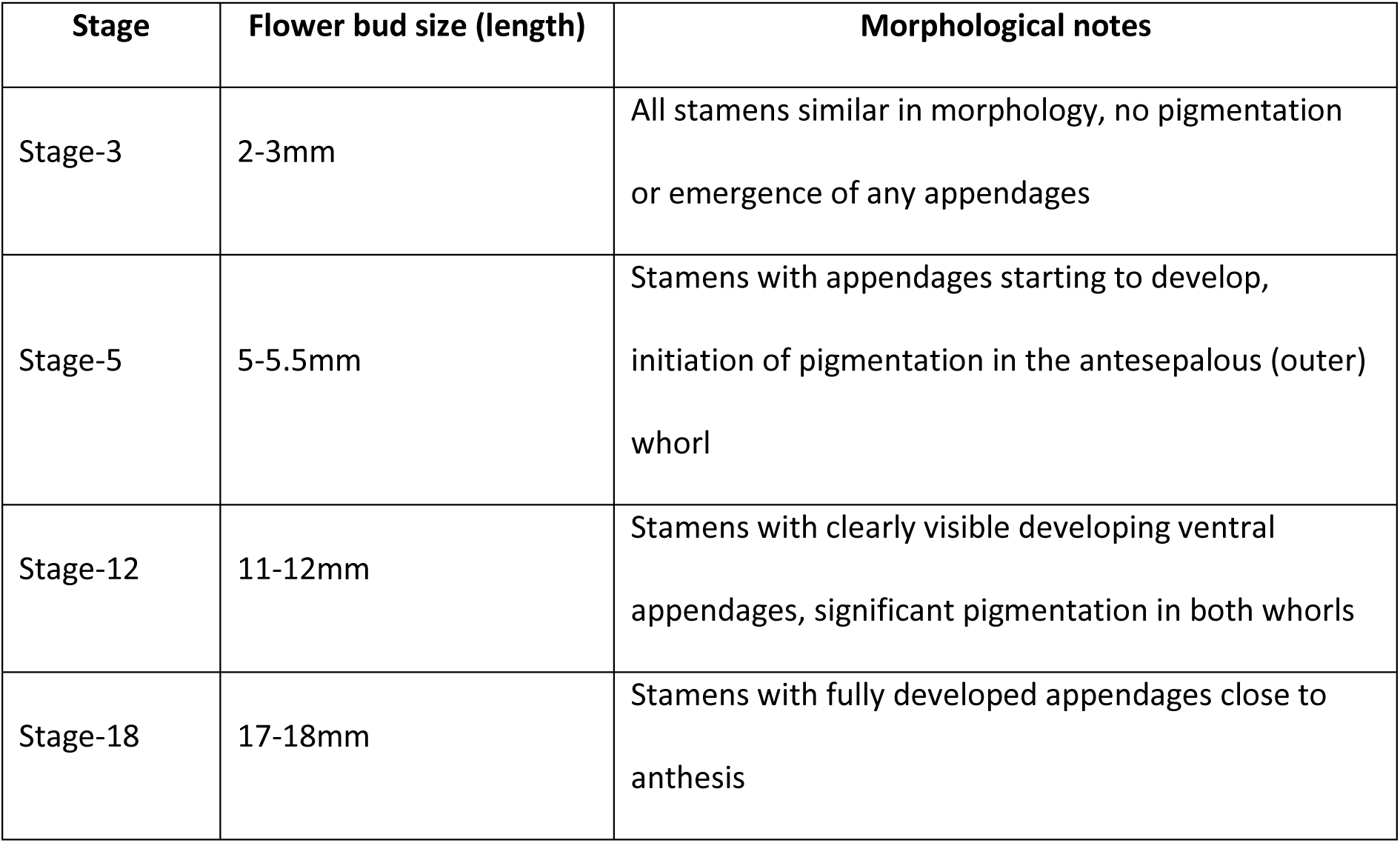
Summary of stages chosen for transcriptomic comparisons.

### RNA isolation from *Arthrostemma ciliatum* stamen

The fresh weight of 4 stamens (one whorl) for the selected developmental stages varied from 0.074mg to∼5mg, so different sampling methods and kits were employed to obtain RNA of good quality and quantity from different stages. Flower buds of appropriate size were collected in moist containers from plants in the greenhouses of the Botanical Garden Munich-Nymphenburg and brought to a nearby dissecting station. Stamens of each whorl were carefully dissected and separated using a microscope and collected in pre-chilled Eppendorf tubes kept in a metal rack on dry ice. This technique allowed for collection of stamen from multiple flower buds without compromising the quality of the RNA and was paramount to get good quantities of RNA from the smaller developmental stages. After complete sampling, the tubes were immediately frozen in liquid Nitrogen for transport to the RNA-lab. Stamens were homogenized in frozen Eppendorf tubes (on dry ice) using plastic pestles, Lysis buffer was added later and RNA was extracted using different kits for different developmental stages. Stage-18 and stage-12 stamen were extracted using Qiagen’s RNeasy® Plant Mini Kit as per the manufacturer’s protocol. Stage-3 and stage-5 stamens were extracted using Qiagen’s RNeasy® Plus Micro kit according to the manufacturer’s protocol. The quality and quantity of the extracted RNA was checked using an RNA ScreenTape® assay in the Agilent TapeStation 4150. Peaks were set and results were visualised using TapeStation Analysis Software 4.1.1 (© Agilent Technologies, Inc. 2021). Summary results of the RNA extractions and the sample names are shown in Appendix-1.

### Sequencing and data analysis

The isolated RNA was diluted with RNase free water to match sequencing requirements and sent to Novogene UK for library preparation and illumina sequencing (RNAseq) with a sequencing depth of 9G. The raw sequence data are available at EMBL accession no #####. All data analysis steps up to and including transcript abundance estimation were performed on a Linux based workstation (Ubuntu 22.04.3 LTS, GNU/Linux 6.5.0-14-generic x86_64).

### Pre-assembly quality control, de-novo transcriptome assembly and post assembly quality control

Raw sequence data was quality checked using FastQC v0.12.1 [40] and summarised with MultiQC v1.17 [41]. Sequencing errors were corrected using Rcorrector v1.0.6 [42] and uncorrectable read pairs were removed using a script obtained from the Transcriptome Assembly Tools github page [43]. Adaptor and quality trimming was performed on the corrected read files using Trim Galore v0.6.10 [44, 45] keeping a minimum output length of 36 bp. Quality checks were performed on the trimmed read files by running FastQC within Trim Galore and later MultiQC.

After the trimming step, reads originating from ribosomal RNA were filtered out by aligning the trimmed read files to the SILVA rRNA database (LSUParc and SSUParc)[46] using Bowtie2 v2.5.2 [47]. The reads that did not align to the SILVA rRNA database were used as the input for de-novo transcriptome assembly using Trinity v2.15.1 [48]. The assembled transcriptome was assessed for completeness using BUSCO v5.5.0 [49]. Read support was measured by mapping/aligning the filtered read files to the assembled transcriptome using Bowtie2 v2.5.2.

### Assembly thinning and abundance estimation

A clustering step using CD-HIT v4.8.1 [50, 51] was performed to reduce sequence redundancy in the assembled transcriptome. Clustering was done using the CD-HIT-EST command with sequence identity thresholds of 98%, 90%, 86% and 80%. Assembly completeness was assessed after clustering for each clustering threshold using BUSCO. The resulting clustered assemblies were used to predict coding sequences using TransDecoder v5.7.1 [52]. Long ORFs (Open Reading Frames) were extracted first and then a blast search using BLAST+ v2.14.1 [53, 54] was done on the extracted long ORFs against the UniProtKB Swiss-Prot database [55] to retain ORFs with homology to known proteins. Then coding regions were predicted using the TransDecoder.Predict command in TransDecoder and any proteins with blast hits were retained.

The clustered transcriptomes were further thinned by subsetting using seqtk v1.4 [56] to keep only those transcripts with coding regions as predicted by TransDecoder. The resulting thinned assemblies were assessed again for completeness using BUSCO. The most complete assembly resulting from these thinning steps (i.e. the 98%-clustered-CDS-predicted assembly) was chosen to estimate transcript abundances using Salmon v1.10.2 [57].

### Differential expression analysis and annotation of transcripts

Differential expression analysis was performed at the transcript level using DESeq2 v1.42.0 [58] in R v4.3.2 [59] on a Windows system following methods adapted from different tutorials and scripts [60–62]. Expression levels were compared in DESeq between the two stamen whorls at each developmental stage. The antepetalous stamen (inner whorl - without pedoconnective; referred to as ‘I’ throughout this analysis) were always used as reference level for these comparisons. Comparisons were also made between adjacent developmental stages for each stamen whorl with the smaller developmental stage as the reference level. A false discovery rate (FDR, alternatively known as the p-adjusted value) threshold of 5% and an absolute log2 fold change (LFC) threshold of 1 were used to identify differentially expressed transcripts (DETs). Sequences of the differentially expressed transcripts from each comparison were extracted and functionally annotated using the online tool Mercator4 [63].

### Pollen counts

Before assessing the pollen grain numbers in antesepalous (outer whorl) and antepetalous (inner whorl) stamens, we checked for any size differences between them by calculating the volume of sporogenous tissue (i.e., the two thecae). Each theca was approximated as an elliptical cylinder and the volumes were calculated accordingly. We also tested for any differences in size between the two thecae within the same stamen. Measurements were taken for all stamen (eight per flower - four antesepalous and antepetalous each) from 10 different pre-anthetic flowers (buds in the size range of 23-25 mm i.e. 1 day before anthesis, recognized by pink colouration of the corolla) to check for size differences. Our results showed that only the length of the thecae was different between the two stamen types (see results section). Hence we compared pollen numbers per anther length to account for these length differences between the stamen types.

To count the number of pollen per anther, one stamen from each whorl was used from 31 pre-anthetic flower buds (one day before anthesis), in order to prevent any pollen loss that might occur after pore opening. The length of the thecae was measured, and the anthers were crushed with a plastic pestle in 1.5mL Eppendorf tubes filled with 100 µL of distilled water. Pollen was counted in a Neubauer Improved Counting Chamber under a Leica DM microscope in 3 replicates for each stamen at 100x magnification. Pictures of pollen from some of these buds were taken at the same magnification and the diameters of the pollen were measured using ImageJ [64].

### Pollen germination experiments

Germination experiments were carried out using a liquid medium prepared according to the recipe for “wet stigmatic plants” of Tushabe & Rosbakh [65], based on the Brewbaker and Kwack’s medium [66]. As a source for the required cations we used MgCl2.6H2O, KNO3 and CaCl2. The pH adjustment was done with NaOH. One stamen of each whorl from an anthetic flower was crushed in 100 µL of the germination medium in 1.5mL Eppendorf tubes and incubated for 30 minutes at room temperature. The tubes were shaken and 10 µL of the solution was transferred onto a microscopic slide, germinating and ungerminated pollen were counted at 100x magnification. This was repeated five times for each stamen type to calculate the pollen viability as percentage of pollen germination. The mean germination percentage per flower and per stamen type was then used for comparisons between stamen whorls.

### Statistical analyses

All statistical analyses on stamen size, pollen size, number and viability for the two stamen types (antesepalous and antepetalous) were conducted using R version 4.5.1 [67]. The lmerTest package [68] was used to fit linear mixed models. Stamen size, pollen size and number were modelled as a function of stamen type as fixed variable with random intercepts for flower or bud to account for repeated measurements: size ∼ stamenType + (1|flower). Models were fit using restricted maximum likelihood (REML) and p-values for fixed effects were calculated using the Satterthwaite approximation for degrees of freedom. The package emmeans [69] was used to check for differences between stamen whorls. For the germination rate, a paired t-test was conducted, as we worked with the mean percentage per flower and per stamen type.

## Results

### Flower development in *Arthrostemma ciliatum* and selection of stages for comparative transcriptomics

In *Arthrostemma ciliatum* sepals are initiated first, followed by the petals, gynoecium and finally the stamens. Before development of the sepals the floral bud is enclosed in a pair of bracts. The gynoecium wall appears free from the hypanthium during young stages. The stamen primordia are inserted on the rim of the hypanthium perpendicular to the floral axis. At the smallest stages observed, antesepalous stamen primordia were always very slightly bigger than the antepetalous ones suggesting that the antesepalous primordia are initiated first. Early in development the stamen becomes folded at the distal part of the filament with the anther apex pointing downwards and inserted between the hypanthium and the ovary wall. The growth of the ovary and stamens is linked with the elongation of the hypanthium. The connective of both stamen whorls grows and thickens dorsally before any elongation of the filament while the ovary wall now appears partly fused with the hypanthium. After some thickening, the connectives of the antesepalous stamens gain a slight pink pigmentation. Thereafter, the connectives of the antesepalous stamen grow bigger than the antepetalous stamens and around this stage the first signs of a ventral outgrowth start to appear on the antesepalous stamens. All stamens grow longer as the petal length increases and a basal increase in connective growth of the antesepalous stamens (pedoconnective development) leads to a strong difference in length between the two stamen whorls. At this point a small ventral outgrowth in the smaller, still unpigmented antepetalous stamens is seen. Subsequently, the appendage and connective growth gains momentum and there is a strong dimorphism between the stamens of the two whorls. Only later does pink pigmentation arise in the antepetalous stamens. Later, filaments start to elongate as the petals grow bigger. The connective near the apex of the antepetalous anthers and the ventral appendage of the antesepalous stamens gain yellow pigmentation at the same time. It is noteworthy that at this point the antesepalous stamens are folded inward at the pedoconnective whereas the antepetalous stamens appear folded at the filament. Around two days before flower opening, the filaments of the longer antesepalous stamen start to twist. This twisting (presumably caused by differential cell division or elongation on one side of the filament) is towards the direction of gravity and moves the stamen in bud, starting a slightly zygomorphic appearance already. During flower opening the stamens all fold out from inside the hypanthium. The twisting movement of the filaments culminates in all the stamens assuming a strongly zygomorphic position, bent downwards with the pores more or less facing the sky (Figure 1: dimorphic). In an anthetic flower the antepetalous stamens are shorter, without a pedoconnective, with a pink filament and a trilobed pink ventral connective appendage (the middle lobe being much smaller than the two lateral); the anther shows a strong curvature forming folds in the theca walls and a yellow colouration of the connective towards the apex (Figure 2). The antesepalous stamen however is longer due to a prominent pink pedoconnective and has an elongated ventral appendage, also trilobed (the lobes much less pronounced than the antepetalous) but with yellow colouration towards the apex; the anther is curved but much less so than the antepetalous stamen (Figure 2). The yellow apices of the antepetalous anthers are bunched together with the yellow ventral appendages of the antesepalous stamen during anthesis attracting bees (Figure 1 dimorphic). Images of the buds and stamens throughout the developmental series can be seen in Appendix-2.

**Figure 2:**
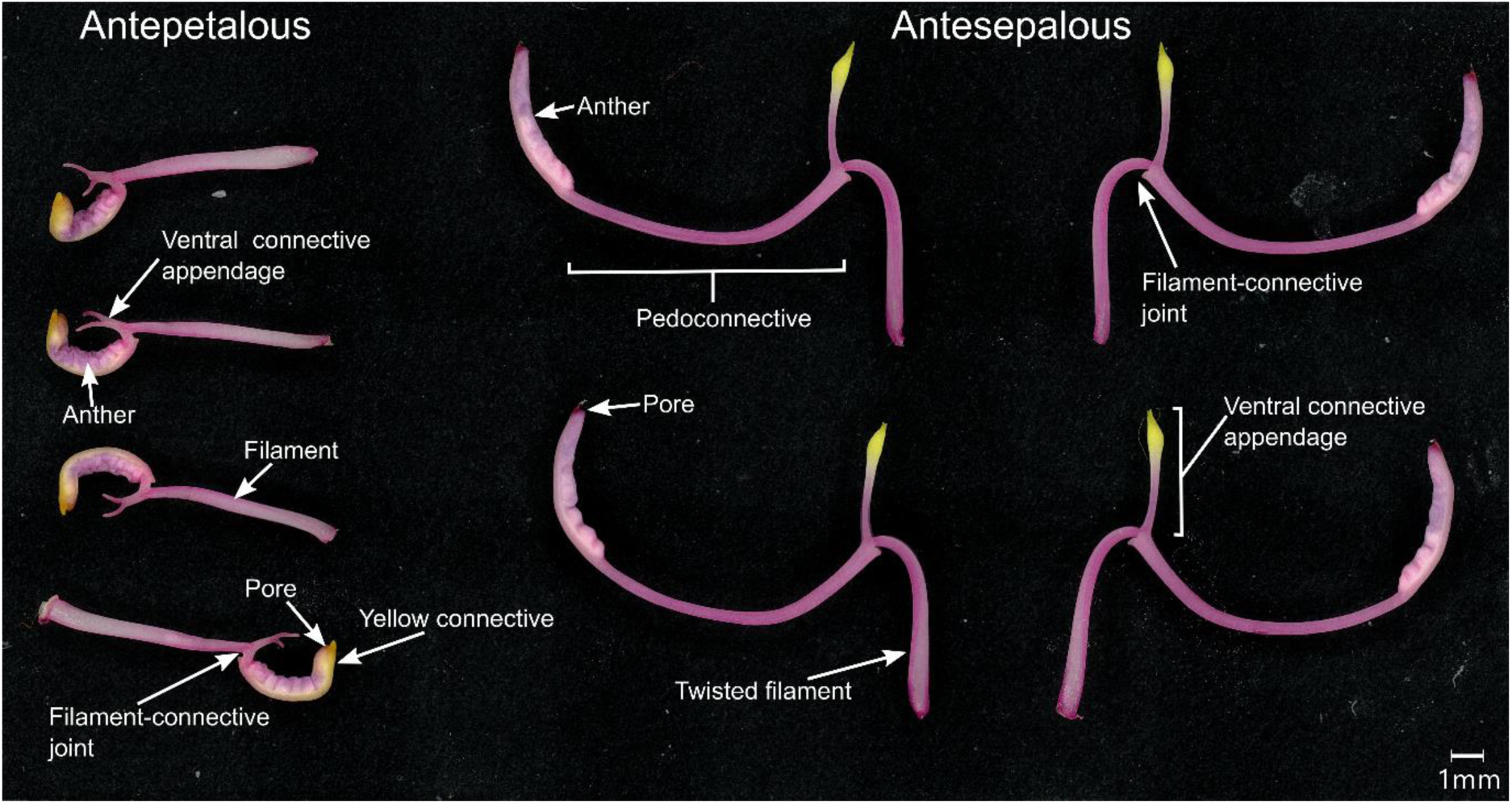
Morphology of mature stamens dissected from an open A. ciliatum flower. Antepetalous stamens (inner whorl) on the left and antesepalous (outer whorl) on the right. Note the size difference and stark morphological differences between whorls. Different parts are labelled in the image.

From these developmental observations four distinct stages of the stamens were chosen for transcriptome comparisons (Figure 3). They were identified based on the morphological characters stated here and the size of the flower buds (see Table 1). Stage-3: All stamens similar in morphology, no pigmentation or sign of any appendages (Figure 3A); 2) Stage-5: Stamens with appendages barely starting to develop (indicated by start of pigmentation in the antesepalous whorl) (Figure 3B); 3) Stage-12: Stamens with clearly visible ventral appendages in the middle of development (Figure 3C); 4) Stage-18: Stamens with fully developed appendages very close to their final form (Figure 3D). Total RNA from entire stamens at these developmental stages was extracted. The antesepalous and antepetalous whorls were separated before extraction. This way we aimed to compare the differences in gene expression between the two stamen whorls at different stages.

**Figure 3:**
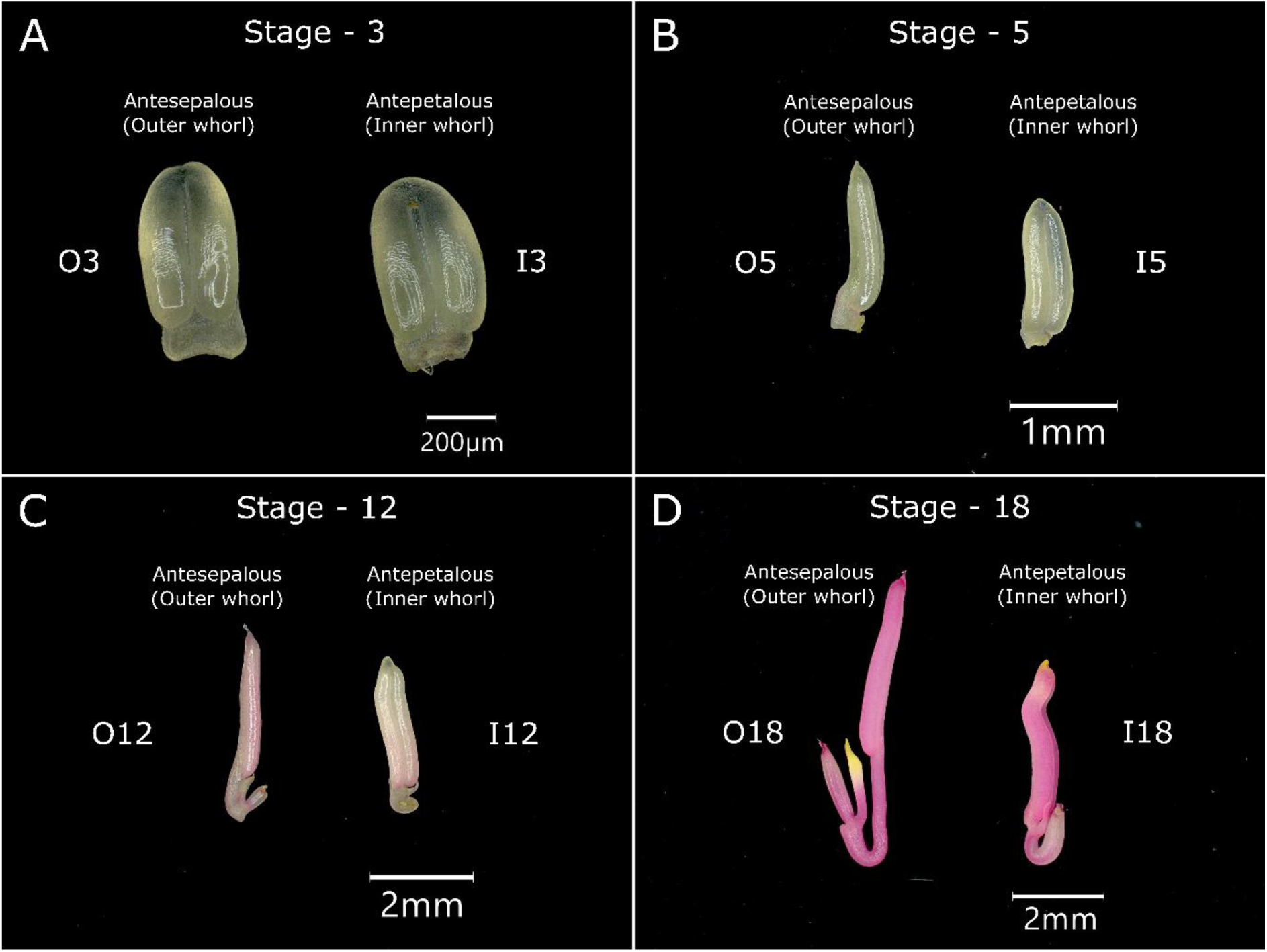
Developmental stages of Arthrostemma ciliatum stamen chosen for transcriptome comparisons. Antesepalous stamen (outer whorl) on the left and antepetalous stamen (inner whorl) on the right for each developmental stage. A) Stage – 3: Flower bud size 2-3mm, no observable morphological difference between the two stamen whorls. B) Stage – 5: Flower bud size 4.5-5.5mm, antesepalous stamen showing initiation of connective appendage. C) Stage – 12: Flower bud size 11-12mm, difference in morphology between the two stamen whorls clearly visible, pedoconnective on antesepalous stamen elongating and the connective appendages on both stamen starting to differentiate. D) Stage – 18 Flower bud size 17.5-18.5mm, both stamen whorls morphologically fully differentiated and nearly ready for anthesis. The abbreviated names for each whorl at each stage are mentioned next to the stamen image (e.g. O3, I3 etc)

### Sequencing and Assembly statistics

Inspection of the MultiQC report for the paired raw reads indicated that all samples passed the quality checks. The number of single reads per sample ranged between 28.7-71.1 million and duplication levels were between 49.7-66.6% (Suppl. Material Appendix-3). After trimming, the number of single reads ranged between 27.8-69.0 million per sample and the adapter content was reduced to zero (Suppl. Material Appendix-3). After removing rRNA contamination the number of single reads ranged between 19.6-60.2 million per sample with duplication levels between 40.5-62.5% (Suppl. Material Appendix-3).

The de-novo transcriptome assembly generated by Trinity using these reads contained 356,512 sequences (transcripts) with an average transcript length of 1161 bases (bp). The length range of transcripts was 185 bp to 28,437 bp. The assembled transcriptome had a read support of 98.41% with 91.58% of the reads aligning one or more times concordantly to the assembly. Assessment of the transcriptome using BUSCO showed that it had a BUSCO completeness of 94.8% (BUSCO notation: C:94.8%[S:3.4%,D:91.4%],F:3.4%,M:1.8%,n:1614). After assembly thinning steps (CD_HIT clustering 98% identity followed by CDS prediction using Transdecoder) the transcriptome contained 130,866 transcripts and had a BUSCO completeness of 94.5% (C:94.5%[S:5.6%,D:88.9%],F:3.5%,M:2.0%,n:1614) This transcriptome was used for transcript abundance quantification with Salmon [57] and is available at EMBL accession no ######.

### Data exploration and differential expression analyses

Of the different data transformation methods used, VST (variance stabilizing transformation) showed better variance stabilisation than rlog (regularised-logarithm transformation) and log2(x + 1) transformation (Appendix-4). Thus, VST data was used to calculate the Euclidean distances between samples in order to assess the similarity between samples. The sample distance heatmap (Figure 4A) shows tight clustering of samples from each developmental stage but the two stamen whorls within each developmental stage do not cluster together. Instead, both stamen whorls from a single sampling event show close relationships in most cases. The PCA and MDS plots also show clustering of samples according to their developmental stages although separation between clusters is clearer in the MDS plot (Figure 4B&C). The PCA covers ∼80% variance in the first two axes. Samples from different developmental stages are clearly separated from each other indicating considerable differences between them. No clustering of stamen whorls was evident within each developmental stage.

**Figure 4:**
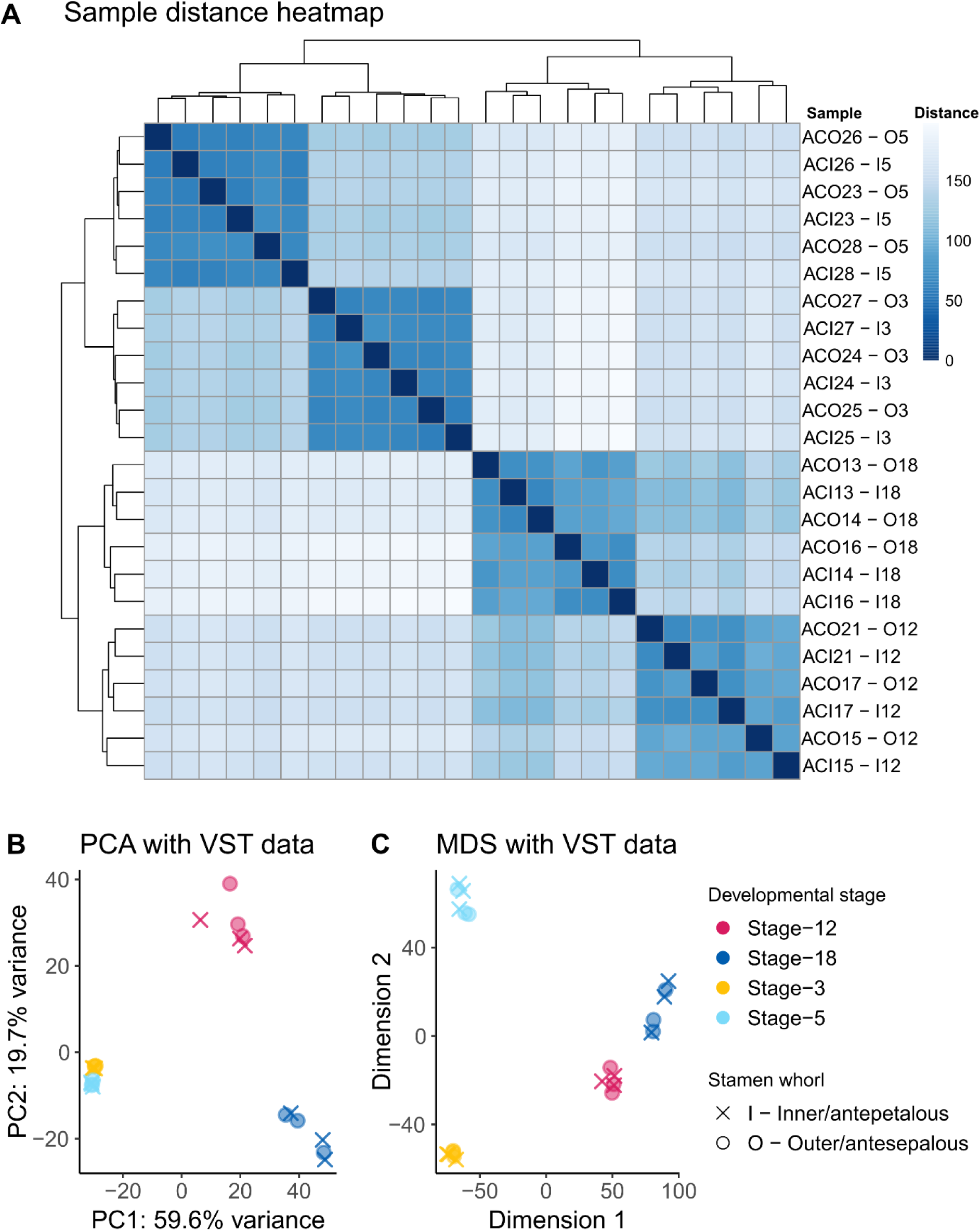
Sample distance comparisons using VST (variance stabilization transformed) data. A) Sample distance heatmap showing Euclidean distances between samples. Sample names and condition are indicated on the right (see Appendix 1 for sample naming convention used in this study). B) PCA plot with VST data; x-axis covers 59.6% variance and y-axis covers 19.7% variance. C) MDS plot using sample distances calculated from the VST data. The points in B & C are coloured by developmental stage and shapes indicate stamen type as shown in legend.

Comparative analyses were conducted within and between the antesepalous (outer-O) and antepetalous (inner-I) stamen whorls at different developmental stages. The inner whorl was used as reference level in all comparisons. Differential expression analyses to identify differentially expressed transcripts (DETs) with an FDR (False discovery rate or p-adjusted value) less than 5% and an absolute LFC >1 yielded 393 DETs between the two stamen whorls at stage-3 (O3 vs I3). Using the same criteria for comparisons between stamen whorls resulted in 277 DETs at stage-5 (O5 vs I5), 281 DETs at stage-12 (O12 vs I12) and 287 DETs at stage-18. The number of up or downregulated transcripts per comparison between stamen whorls were relatively similar (Figure 5). However, the comparisons between adjacent developmental stages of each stamen whorl showed a much higher number of DETs using the same FDR and LFC thresholds. In the antepetalous (inner) whorl, comparison between stage-3 and stage-5 stamens (I5 vs I3) gave 5403 DETs. 8310 DETs were identified when comparing stage-5 and stage-12 (I12 vs I5) whereas between stage-12 and stage-18 there were 2371 DETs. Similarly, comparisons between stages in the antesepalous (outer) whorl resulted in identification of 4641 DETs between stage-3 and stage-5 (O3 vs O5); 7409 DETs between stage-5 and stage-12 (O5 vs O12); and 2608 DETs between stage-12 and stage-18 (O18 vs O12). For both stamen whorls, a large difference between the number of upregulated and downregulated genes when comparing stage-3 with stage-5 is evident (Figure 5).

**Figure 5:**
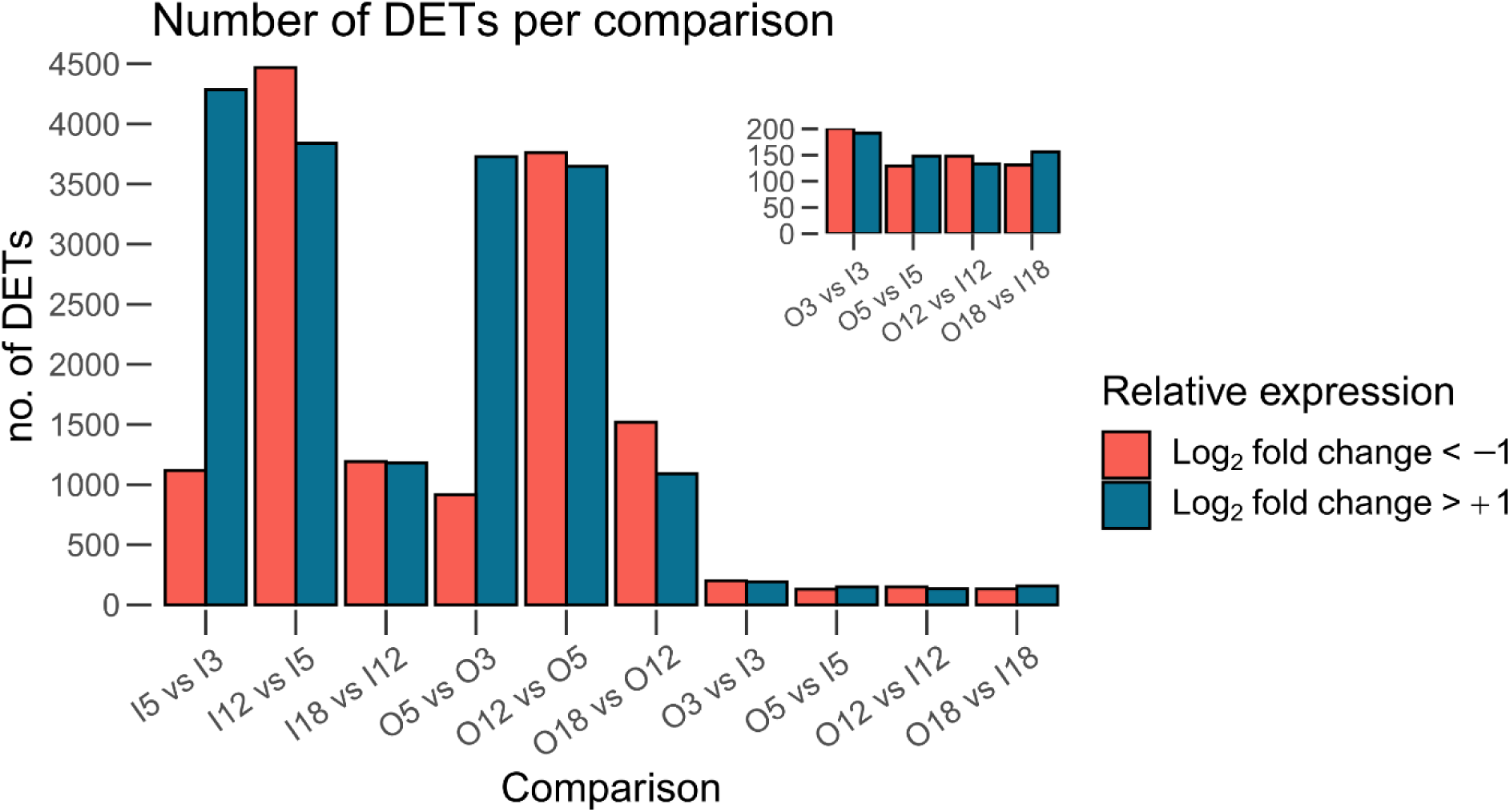
Number of differentially expressed transcripts per comparison. Bar plot showing number of upregulated (LFC >+1, blue) and downregulated (LFC<-1, salmon) transcripts with an adjusted P value <0.05 for each inter-developmental stage comparison and inter-whorl comparison. The letters in the comparison stand for the stamen whorl (I - Inner whorl/antepetalous, O - outer whorl/antesepalous) while the numbers refer to the developmental stage (e.g. I5 vs I3 refers to the compares the inner whorl stamens at stage-5 and stage-3 while O3 vs I3 refers to the comparison of outer and inner stamen whorls at stage- 3) The right-hand-side condition in each comparison shown on the y-axis (e.g. I3 in I5 vs I3) was always used as the reference level. Inset shows the inter-whorl comparisons (column pair 7-10) magnified.

### Annotation of DETs and identification of transcripts involved in stamen development

The DETs from the inter-whorl comparisons (O vs I) were annotated by Prot-scriber and Swissprot annotations in Mercator4. The annotated transcripts involved in stamen development and elongation were identified based on extensive literature searches and separated out as transcripts of interest (see suppl. Material Appendix-5 for merged results tables of all the genes of interest from the inter-whorl comparisons). The count data for these transcripts of interest was checked for reliability.

All the DETs summarized in the following paragraph have been previously shown to be involved in stamen development or determining filament length in *Arabidopsis* and other plant species. Several transcripts involved in phytohormone action, stamen development and elongation were observed in all the inter-whorl comparisons (Figure 6, Appendix-5). More specifically, in the comparison O3 vs I3 an auxin influx transporter (AUX/LAX), a cytokinin dehydrogenase (CKX) and multiple MYB class R2R3 transcription factors were upregulated whereas transcripts coding for an EPF/EPFL precursor polypeptide and a DELLA protein were downregulated in O3 stamen (Figure 6A). The comparison O5 vs I5 showed downregulation of transcripts coding for Auxin efflux transporter family (PIN-LIKES), Auxin binding protein 1 (ABP1), EPF/EPFL precursor polypeptide and a WOX type transcription factor while a JAZ-TPR transcriptional repressor complex protein was upregulated in condition O5 (Figure 6B). Transcripts of SCL (Scarecrow-like protein) and Agamous-like MADS-box transcription factor (MADS/AGL TF) were also differentially expressed in this comparison. In condition O12 a gibberellin receptor (GID1), a phosphoinositide phosphatase (SAC1), a cysteine proteinase involved in programmed cell death (VPE) and an autophagy related protein (ATG 16) were downregulated compared to I12; on the other hand an AP2/ERF (ethylene response factor) family transcription factor, an S-LOCUS EARLY FLOWERING 3 (ELF3) homolog, a Brassinosteroid signalling kinase (BSK7) and a cytokinin phosphoribohydrolase (LOG) were upregulated in O12 compared to I12 (Figure 6C). Stamens of condition O18 showed upregulation of a Phytochrome (PHY-B), a LOB (lateral organ boundary) domain containing protein (AS2/LOB transcription factor) and a WAV3 family E3 ubiquitin ligase protein compared to condition I18. An autophagy related protein (ATG 16), a MADS-Box transcriptional regulator (AP1), a Histone trimethyltransferase (ATX1) and RADIALIS transcription factors were all downregulated in O18 stamen compared to I18 (Figure 6D). The LFCs and p-adjusted values of some relevant DETs are shown in Appendix-5 along with their Mercator4 annotations.

**Figure 6:**
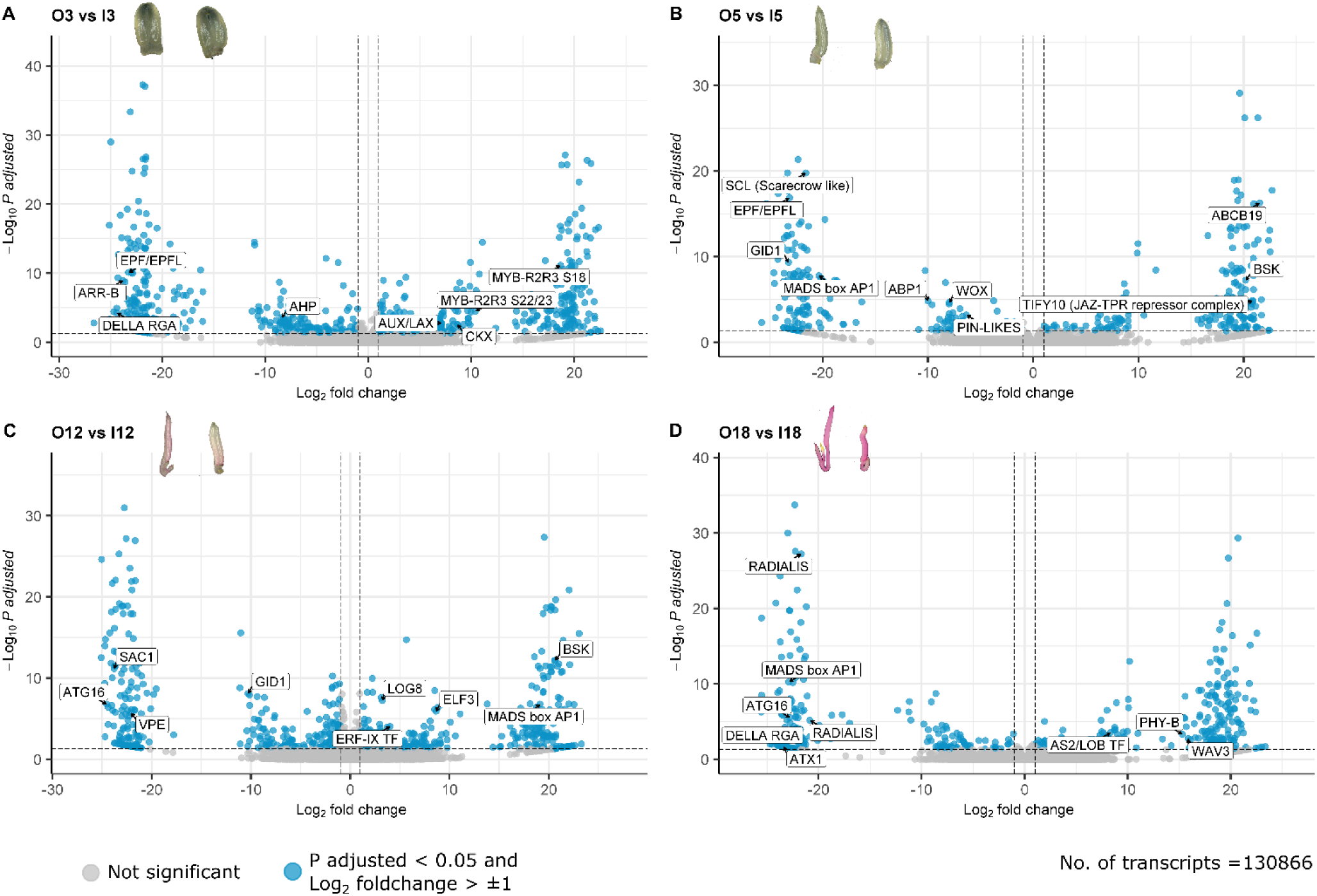
Volcano plots for each inter whorl comparison. A) O3 vs I3: Outer whorl at stage 3 compared to inner whorl at stage 3. B) O5 vs I5: Outer whorl at stage 5 compared to inner whorl at stage 5. C) O12 vs I12: Outer whorl at stage 12 compared to inner whorl at stage 12. D) O18vs I18: Outer whorl at stage 18 compared to inner whorl at stage 18. Points in blue are differentially expressed transcripts with a p adjusted value of less than 0.05 and a log2 fold change of more than one on either side of zero. The gene names of the transcripts of interest as annotated by Mercator4 are marked on the plots. The inner whorl (e.g. I3 in O3 vs I3) was always used as the reference level for the comparisons so the log2 fold changes are of the outer whorl stamen with respect to the inner whorl stamen.

### Theca size, pollen quantity and quality in *Arthrostemma ciliatum* stamen types

Statistical tests using Linear mixed models (LMM) and post hoc comparisons using the emmeans (Estimated marginal means) package showed no significant difference in theca size within each stamen. There was no significant difference between the cross-sections of antesepalous (AS) and antepetalous (AP) thecae (emmeans contrast AP-AS estimate= -0.00393mm^2^, Std. err. (SE)= 0.00387, degrees of freedom (df)= 69, t ratio= -1.015, p value = 0.3136) (Figure 7B) while there was indeed a significant difference in theca lengths (AP emmean= 4.26mm, 95%CI [4.15, 4.37]; AS emmean=5.26mm, 95%CI [5.15,5.37]; SE= 0.0481, df=10.4), with antesepalous stamens being an estimated ∼0.99mm longer than the antepetalous ones (SE= 0.0259, df= 69, t ratio= -38.625, p value <0.0001) (Figure 7A & Appendix-6). This led to the conclusion that the difference in volumes between the two stamen types (Figure 7C & Appendix-6) (emmeans contrast AP-AS estimate= -0.119mm^3^, SE=0.0184, df= 69, t ratio= -6.463, p value <0.0001) is the consequence of theca length. So, pollen number per length of anther was calculated to correct for the observed differences in length between the two stamen types. We found that the counted pollen per mm length of theca followed a bimodal distribution related to date of sampling. The variation in the number of pollen between flower buds was the largest (Std. Dev. = 1107.7), where the date of sampling had a big influence (Appendix-7). Analyses showed a significant difference in pollen per mm between stamen types (emmeans contrast AP-AS estimate= 770, SE= 151, df= 30, t ratio = 5.100, p value < 0.0001), with antepetalous stamen having an estimated marginal mean of 3702 pollen/mm (95 % CI [3245, 4159], SE = 226, df = 37.4), and antesepalous stamen having an estimated marginal mean of 2932 pollen/mm (95 % CI [2475, 3389], SE = 226, df = 37.4) (Figure 7D). Comparing the average pollen diameters from antesepalous and antepetalous stamen using LMM showed a significant difference in size (emmeans contrast AP-AS estimate= -3.21 µm, SE= 0.121, df= 339, t ratio= -26.541, p value <0.0001) with antesepalous pollen (AS) having an estimated mean diameter of 33.3 µm (95 % CI [32.7, 33.8], SE = 0.241, df= 5.69), and antepetalous (AP) pollen being smaller with an estimated mean diameter of 30.0 µm (95%CI [29.4, 30.6], SE= 0.241, df= 5.69) (Figure 7E & Appendix-8).

**Figure 7:**
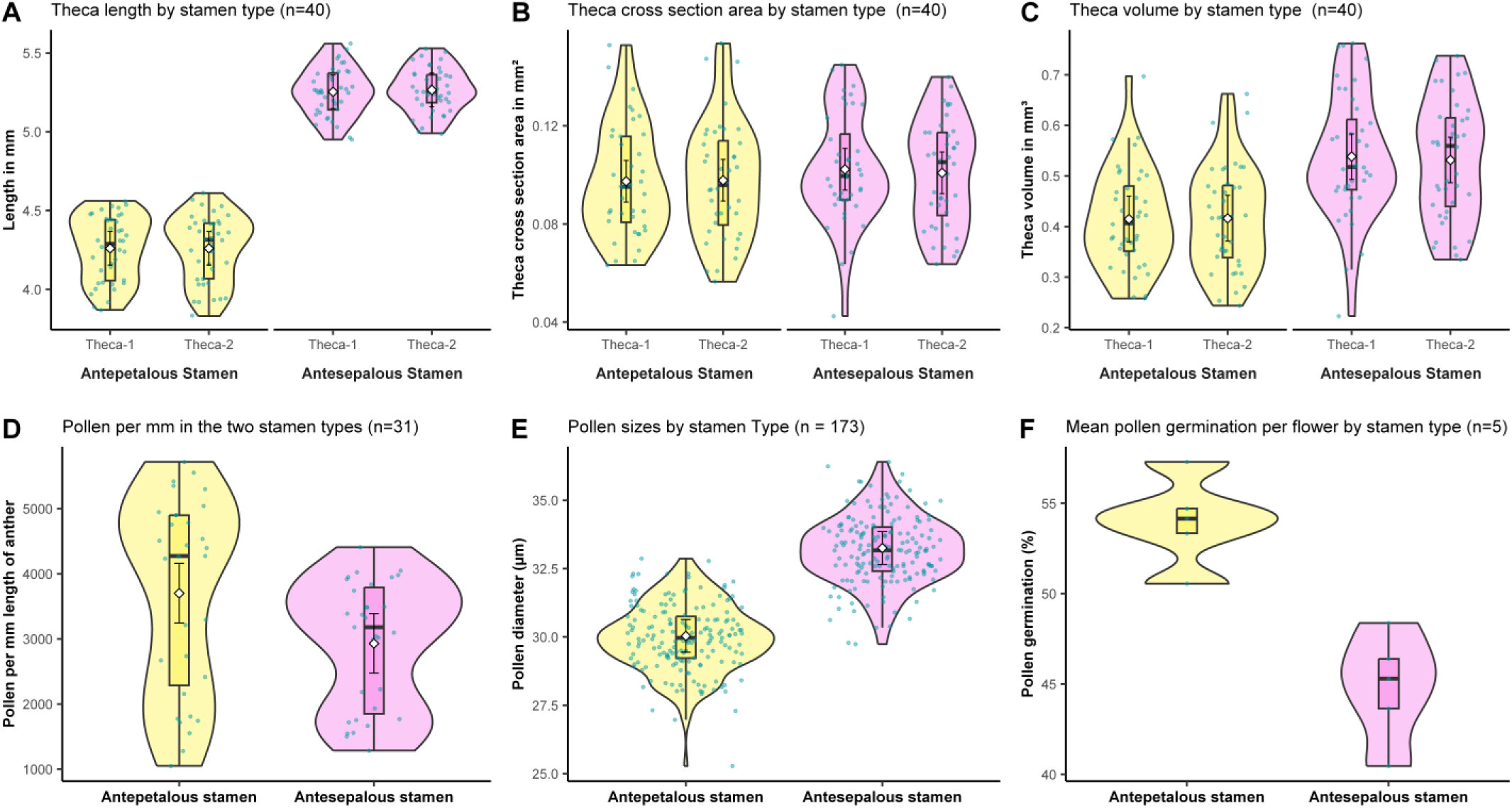
Stamen size, pollen number, size and viability comparisons. Violin-box-jitter plots comparing stamen size, pollen number, size and pollen viability between the two stamen types (antesepalous-outer whorl; antepetalous-inner whorl). The estimated marginal means (white rhombi) and 95% confidence intervals (black stapled lines) obtained via emmeans package are shown overlapping the boxplot where available. The sample sizes mentioned are biological replicates. A) Theca length in mm for the two stamen types. B) Theca cross section area in mm² for the two stamen types. C) Theca volume for the two stamen types. Theca volume was calculated by multiplying the length with the cross section area. D) Pollen per mm length of the thecae for the two stamen types. E) Pollen sizes for the two stamen types. F) Mean pollen germination per flower for the two stamen types (emmeans not run on this dataset).

The viability of pollen from antepetalous stamen anthers was significantly higher than those from antesepalous stamen anthers (paired t-test, t =4.2201, df= 4, p value = 0.01348, 95% CI [3.137944, 15.207867) (Figure 7F & Appendix-9). Approximately 45% of pollen extracted from antesepalous stamen anthers were able to germinate. Whereas, the percentage of germination was substantially higher in pollen from antepetalous stamens with ∼54%.

## Discussion

Stamen dimorphism (heteranthery) is a widely distributed phenomenon among flowering plants but it is especially prevalent in certain buzz pollinated groups like some Fabaceae, Solanaceae and Melastomataceae among which melastomes represent the largest group [33]. Ecological aspects of heteranthery have been extensively studied as detailed in the introduction, yet there has been no prior study focusing on the genetic basis underlying stamen dimorphism in Melastomataceae. This study marks the first detailed foray into this field for this family, apart from a study on the genome of *Melastoma dodecandrum* Lour. [38], which gives little detail on genes involved in heteranthery. There have however been similar studies conducted on stamen dimorphism in *Cassia biscapsularis* L. and *Monochoria elata* Ridl. where they identified several gene families that possibly cause stamen dimorphism [37, 39]. We aimed to build on these studies with respect to the genes that might be responsible for heteranthery by comparing the gene expression profiles of the heterantherous stamen whorls of *Arthrostemma ciliatum* at different developmental stages and have gained promising first insights as discussed below.

### Stamen development in *Arthrostemma ciliatum*

The order of initiation of organ primordia in *Arthrostemma ciliatum* is similar to the pattern seen in previously studied melastomes [20, 70–72]. Stamen are the last floral organs to be initiated as is the general trend in Melastomataceae [71]. The order of initiation of stamen whorls (outer antesepalous followed by inner antepetalous stamen) is correlated to their final size (antesepalous stamen larger than antepetalous). The concept of ‘imprinted shape’ suggested by previous studies [20, 73] seems relevant here as the shape of connective appendages mirrors the space available during their development. The size of anthers during development is constrained by the space available for growth between the hypanthium and the ovary wall. A large part of the hypanthium of our study species remains unfused with the ovary wall and elongates during development, forming extraovarian chambers into which the anthers grow. This supports the hypothesis that a longer perigynous hypanthium favours an elongation of the anthers and stamen which might then also allow for the development of a pedoconnective and appendages [20].

Folded stamen in bud are typical of all Melastomataceae and many other families in Myrtales [74]. In *A. ciliatum* a strong curvature of the poricidal anthers is observed at anthesis after the stamens unfold out of the hypanthium (Figure 2). This curvature is more pronounced in the antepetalous/inner whorl and causes the walls of the thecae to attain partial transverse folds which might serve as a pollen dosing mechanism [75]. If true, this mechanism would mean that the pollen in the antepetalous stamen is released over multiple pollinator visits, increasing the chance of successful pollination. Further developmental studies at a cellular level are needed in the future to understand whether the main cause of pedoconnective elongation in the antesepalous stamens is cellular division or elongation.

### Candidate genes responsible for staminal dimorphism in *Arthrostemma ciliatum*

Exploratory analyses using the count data showed that the differences among samples was mainly between developmental stages, where samples from the same stage shared a similar expression profile. The two stamen whorls originating from the same sampling event showed minimal differences within each developmental stage. This indicates that the expression profiles of antesepalous and antepetalous stamen within individual flowers are very similar. We did not observe clustering of samples according to stamen whorl which pointed towards the same conclusion but it could be that three biological replicates are too little to see consistent differences among whorls at each stage. The same trends were observed from the number of differentially expressed transcripts per comparison. A larger number of DETs in the inter-stage comparisons compared to the inter-whorl comparisons indicates that the expression profiles of the antepetalous and antesepalous stamens at the same developmental stage are very similar. There is a large difference between the number of upregulated and downregulated genes in stage-5 compared to stage-3 in both stamen whorls (Figure 5). This suggests that the increasing morphological complexity from stage-3 to stage-5 is reflected in the upregulation of a large number of genes which are probably inactive in early stages.

R2R3 type MYB transcription factors along with jasmonate signalling components (including JAZ-TPR), DELLA, GID1, ARFs, gibberellin and other phytohormone signalling components have previously been shown to control stamen development and filament elongation in *Arabidopsis* [76–81]. DELLA proteins are known to repress stamen development in *Arabidopsis* [76, 81–83] and they are degraded by the action of GID1+gibberellin via ubiquitination [81]. This promotes jasmonate synthesis and activates the R2R3 type MYB transcription factors which in turn are involved in various stages of stamen development including affecting filament elongation [81]. The differential expression of these genes between the two stamen whorls in *A. ciliatum*, albeit at different developmental stages, potentially indicates that the difference in connective lengths (leading to pedoconnective formation in the antesepalous whorl -O) might be controlled by a mechanism involving the same gene families. Assuming that these components have the same functions in *Arthrostemma,* the upregulation of R2R3 MYB transcription factors and downregulation of DELLA protein in O3 support this hypothesis while the fact that GID1 is downregulated in O12 seems to contradict it.

Other differentially expressed genes which are known to be involved in determining stamen length are the EPF/EPFL family peptides and the S-LOCUS EARLY FLOWERING genes. Mutations in EPF/EPFL family genes have been shown to cause shortened filament phenotypes in *Arabidopsis* and *Triticum* due to reduced filament cell proliferation [84–86]. The EPF/EPFL peptides act as ligands for ERECTA and Somatic Embryogenesis Receptor Kinase (SERK) family receptor complexes which initiates a signal cascade involving MAP (mitogen activated protein) kinases and enhances filament cell proliferation [84]. It is possible that differential expression of one of the EPF/EPFL family genes between the two stamen whorls causes proliferation of cells in the pedoconnective of antesepalous stamen causing its elongation. The S-locus genes on the other hand are involved in self incompatibility [87, 88] and the EARLY FLOWERING (ELF) genes control heterostyly in many plant species [87–91]. Initial indications from observations in the field suggest that there are indeed two style morphs in *A. ciliatum* (S.Kotagal pers. obs.). The isomorphic flowers seem to always have styles which declinate later during anthesis which positions the stigma in a similar space as the pores of the long stamen in the dimorphic flowers.

Heterostyly has not been described in *Arthrostemma ciliatum* so far, but from the above observations, and the fact that the two morphs occur in close geographical proximity, we think it is highly probable that this might be the case in this system. Further studies are underway to verify this phenomenon but the differential expression of ELF homologs between the two stamen types indicates that these genes merit further study. It is possible that the S-locus ELF homologs are involved in controlling heteranthery and heterostyly in this species similar to what is seen in other heterostylous species but affecting connective length instead of filament length.

The upregulation of Vacuolar processing enzyme (VPE) and autophagy related proteins in the antepetalous stamen of stage-12 (I12) suggests that the morphological differences of the antepetalous (inner) stamen compared to the antesepalous (outer) stamen (absence of pedoconnective, smaller connective appendage, highly dissected appendage lobes) (Figure 2) might be caused by increased programmed cell death (PCD) in the inner whorl. It has been established that programmed cell death (otherwise known as apoptosis) plays a major role in morphogenesis in plants and animals [92–94], and the functions of VPE in plant development and stress induced cell death are well studied [95–98]. VPE is a cysteine protease which is functionally similar to caspases and is involved in mediating PCD via vacuole rupture and a proteolytic cascade [95, 98]. It is possible that increased VPE in the inner stamen causes increased PCD which might prevent excessive cell division leading to the reduced size of the connective and elaborate connective structures.

In our developmental studies we observed that the outer stamens move gravitropically in bud (by a twisting growth of the filament) earlier than the inner whorl at around the same developmental stage as stage-18. This is mirrored by the expression patterns of a phytochrome (PHY-B) and WAV3 ubiquitin ligase, which were upregulated in antesepalous (outer whorl) stage-18 stamens. The phytochromes are known to be involved in flower opening [99, 100] while WAV3 ubiquitin ligases have been shown to play roles in gravitropic responses in roots [101]. These genes might be involved in causing the early gravitropic movement of outer stamen leading to the observed staminal asymmetry in bud before anthesis. The differential expression of a RADIALIS transcription factor at this stage suggests that homologs of this gene might also be involved in controlling the symmetry of stamen in this species. The RADIALIS transcription factors are known to control floral symmetry together with DIVARICATA and CYC genes [102, 103] and it is plausible that these genes also control the dorsiventral symmetry of stamen in Melastomes.

In summary, we found that the transcripts of the Jasmonate and Gibberellin signalling pathway (involving DELLA, GID1, ARF and MYB transcription factor homologs), EPF/EPFL family of genes, S-Locus ELF homologs and autophagy related genes (VPE) are differentially expressed between the two stamen types. Homologues of these genes have all previously been found to be involved in stamen development and controlling stamen length. Thus, we believe that these genes are putative candidates causing the length differences among stamen whorls in dimorphic *A. ciliatum* and merit further study. ARFs were also suggested to be the cause of differential length of stamens in the study on *Cassia biscapsularis* [37] while in *Melastoma dodecandrum* SAUR-like homologs were suggested to be involved in causing heteranthery by interacting with auxin [38]. In the study on *Monochoria elata* [39], genes involved in hormone signalling, specifically the Abscisic acid (ABA) and Jasmonate signalling pathways were suggested to cause differential stamen elongation. However, it should be noted that both *Cassia* and *Monochoria* are distant relatives to *Arthrostemma,* occurring in different clades in the Rosids and Monocots respectively. Heteranthery in these two species and in *Arthrostemma* is, strictly speaking, not homologous - in *Cassia and Monochoria* there is no elongation of the connective (pedoconnective) as in *Arthrostemma* and any similarities or differences in the genetic mechanisms among these species need to be corroborated with further studies. Generation of genome level assemblies coupled with further analyses of our dataset using WGCNA and additional studies comparing this dataset to the isomorphic form of *A. ciliatum* and other species of *Arthrostemma* will further help to narrow down the genes involved in causing heteranthery in this group. Techniques for functional validation of these genes such as transformation or knockdowns still need to be developed for melastomes and in general such studies still have much work to do to successfully identify the causal genes for heteranthery. But this study represents the first major stride in this direction for Melastomes and we aim to follow up on the questions brought up here in the near future.

### Heteranthery in *Arthrostemma ciliatum* cannot be clearly linked with the “division of labour” hypothesis

*Arthrostemma ciliatum* at first glance seemed to be a classical example of the division of labour hypothesis. The short antepetalous stamen with yellow-coloured apices (Figure 2) bunched together with the yellow ventral appendages of the longer stamen attracting the pollinators’ attention and providing a reward suggested the function of the antepetalous stamen as the feeding stamen. While the longer antesepalous stamen camouflaged against the petal background and spatially separated from the other whorl due to the elongated pedoconnective, suggested their function as pollinating stamen and that they place pollen on the ‘safe sites’ of the bee body. Our findings however, show that - in this morph of *A. ciliatum* the antepetalous stamens produce smaller pollen in a greater number while also having better germinating potential than the antesepalous stamen. This seems contrary to the theory that antepetalous stamen serve as feeding stamens since previous studies have indicated that the pollinating stamen have smaller, more viable pollen in a greater number than feeding stamen in heterantherous species [104, 105]. It could also be argued that the higher pollen number and smaller size in the antepetalous stamen together with the folded theca walls, act as a pollen dosing mechanism and serve to ensure multiple pollinator visits while the bigger, although less numerous pollen from the antesepalous stamen, contribute to successful pollination as they might be able to produce longer pollen tubes [106, 107]. But the fact that the antesepalous pollen were found to be less viable in this morph of *A. ciliatum* advocates against this theory. The bigger pollen in the longer stamen and smaller pollen in the smaller stamen is similar to the phenomenon found in heterostylous species where the size of pollen corresponds to the size of the style in the opposite morph. There is still considerable debate concerning the relationships between pollen size, number, stigma/style length and pollination strategies (see [105, 108–112] and references within). Data for heterantherous species, especially within Melastomes, is severely lacking in this respect and since these relationships seem to be group specific, it is hard to lean towards one hypothesis or the other at the moment. For us however, the closing argument concerning the division of labour in the dimorphic form of *A. ciliatum* is with the spatial positioning of the stigma relative to the two stamen types (see Figure 1 Dimorphic and page 5 in Appendix-2). The stigma of the heterantherous morph, throughout the life of the anthetic flower, is surrounded by the antepetalous stamen in such close proximity that any removal of pollen by vibrating bees would invariably deposit pollen from the same antepetalous stamen on it. This, coupled with our observation of fruit and seed set upon selfing in the greenhouse, provides a strong case for self-pollination being the main strategy of this morph. Although this does not exclude the presence of incompatibility mechanisms in the wild, it would explain the observed higher viability of the antepetalous pollen.

In summary, we found that antepetalous stamen show both feeding and pollinating function in dimorphic *A. ciliatum*, leading to doubt regarding the function of antesepalous stamen. However, the presence of isomorphic populations of the same species in close geographic proximity suggests the possible existence of a distylous (or at least distyly-like) system. At least some individuals of the isomorphic form of *A. ciliatum* have slightly longer styles that declinate later during anthesis (Pers. obs.; see the two isomorphic flowers in Figure 1 - Isomorphic) making their stigma spatially closer to the pores of the long antesepalous stamen of the dimorphic form, increasing the chances of outcrossing. This putative outcrossing mechanism would explain the consistent co-occurrence of the heterantherous form of this species with the short stamened form. Furthermore, Almeda and Chuang [113] suggested that the two morphs differ in their karyotypes. This poses more questions regarding the breeding system of this species and whether the two morphs constitute an interbreeding population or two distinct reproductively isolated populations needs to be tested via further field studies of the natural populations. A comprehensive phylogeny of the genus and the tribe including several populations of both morphs of *A. ciliatum* across its native range would provide the desperately needed phylogenetic context to better understand the evolution and significance of heteranthery in this group and work towards this is already underway.

## Conclusions

From our developmental observations we confirm that stamen development in the heterantherous form of *Arthrostemma ciliatum* shows similar patterns as seen in other melastome species. Based on differential expression analyses using our developmental framework we identified the Jasmonate and Gibberellin signalling pathway (involving DELLA, GID1, ARF and MYB transcription factor homologs), EPF/EPFL family of genes, S-Locus ELF homologs and autophagy related genes (VPE) as putative candidates that might be causing heteranthery in *A. ciliatum.* By comparing our results with previous studies, we highlight the convergent nature of heteranthery and theorise that distantly related plant groups might have different genetic bases controlling this trait. We also suggest that WAV3 ubiquitin ligase, Phytochrome-B (PHY-B) and RADIALIS transcription factors are putative candidates determining the zygomorphy of the stamens. The transcriptomic dataset we generated provides initial insights into the genetic basis of heteranthery in *A. ciliatum* while also reinforcing the foundation for further evolutionary studies on this phenomenon in Melastomataceae and across angiosperms. Furthermore we show that heteranthery in *A. ciliatum* cannot be explained by the classical division of labour theory as the shorter stamen have more pollen with better viability. We instead suggest the existence of heterostyly in combination with heteranthery as an outcrossing mechanism to explain the occurrence of two staminal and stigma morphs in this species.

## Supporting information

Appendix-1: Sample names and summary table of RNA extractions

Appendix-2: Developmental series images of flower buds and stamen

Appendix-3: MultiQC reports of raw data; after trimming and after removal of rRNA

Appendix-4: Scatterplot of transformed counts from two samples

Appendix-5: DESeq results tables with differentially expressed genes of interest from each comparison showing FDR, LFC, Counts and annotations

Appendix-6: Violin plots of theca size by stamen type and flower

Appendix-7: Violin plots of length of anthers used for pollen counts and pollen numbers by stamen type

Appendix-8: Violin plots of pollen size by stamen type and flower

Appendix-9: Violin plots of pollen germination by stamen types and flower

## Supplementary Data/Additional files

**Appendix-1:** (Appendix-1_RNA_extraction_summary_table.pdf) Sample names and summary table of RNA extractions

**Appendix-2:** (Appendix-2_Development_series_images.pdf) Developmental series images of flower buds and stamen

**Appendix-3:** (Appendix-3_Multiqc_reports\ .html) MultiQC reports of raw data; after trimming and after removal of rRNA

**Appendix-4:** (Appendix-4_scatterplot_count_transformations_noleaf.pdf) Scatterplot of transformed counts from two samples

**Appendix-5:** (Appendix-5_Genes_of_interest_inter-whorl-comp_merged.xlsx) DESeq results tables with differentially expressed genes of interest from each comparison showing FDR, LFC, Counts and annotations

**Appendix-6:** (Appendix-6_Theca_size_comparisons_between_stamen_types_by_flower.png) Violin plots of theca size by stamen type and flower

**Appendix-7:** (Appendix-7_Pollen_number_plots_1.png) Violin plots of length of anthers used for pollen counts and pollen numbers by stamen type

**Appendix-8:** (Appendix-8_Pollensizes_1.png) Violin plots of pollen size by stamen type and flower

**Appendix-9:** (Appendix-9_Pollen_germination_by_StamenType_and_flower.png) Violin plots of pollen germination by stamen types and flower

## Declarations

### Ethics approval and consent to participate

Not applicable

### Consent for publication

Not applicable

### Availability of data and materials

Raw RNASeq data generated during this study will be made available under the accession number xxxx in EMBL. The raw data of stamen size, pollen number, pollen size and pollen viability will be deposited in Zenodo @ doi. All scripts used in this study will be made available at github@Suvrat.

### Author contributions

AS- Anna Schlick; CS- Christian Siadjeu; EYH- Emy Yue Hu; GK- Gudrun Kadereit; SK- Suvrat Kotagal Conceptualization - SK & GK; Methodology – SK, AS, CS, EYH & GK; Formal analysis – SK, AS, CS & EYH; Investigation – SK & AS; Resources – GK; Data curation – SK; Writing-original draft – SK & GK; Writing – review & editing – AS, CS, EYH, SK & GK; Visualization – SK, AS, EYH & CS; Supervision – GK, CS & EYH; Funding acquisition – GK.

## Acknowledgements

We thank the gardeners (in particular Andreas Richter and Harald Loose) at the Botanical Garden Munich-Nymphenburg for growing and maintaining the plants used in this study and for their support during the delicate sampling process. Eva Facher and Seraina Rodewald for their guidance during the developmental studies; Martina Silber, Silvia Wienken, Jean-Baptiste Chazalon and Alina Höwener for their guidance and help with the lab work.

