## Appendix-1: Sample names and summary table of RNA extractions for "Insights into the functional and genetic basis of heteranthery in *Arthrostemma ciliatum* Pav. ex D.Don (Melastomataceae)"

**Table 1:** Sample table for *Arthrostemma ciliatum.* Sample names are indicative of the species (AC), stamen whorl (O-outer/antesepalous and I-inner/antepetalous) and sampling event number. The Condition column for each sample refers to the stamen whorl and the developmental stage at which it was sampled. RINe refers to RNA Integrity Number equivalent output by the Agilent Tape station analysis, a value of 10.0 indicates high RNA integrity and minimal degradation. Conc refers to RNA concentration after isolation obtained by the Agilent Tapestation analysis. VolEl refers to the final volume of elution.

| Sample name | No. of flowers/ stamen | Condition (whorl+ bud size) | Date | Sampling time (24h) | Size of buds | RNA extraction kit | RINe value | RNA Conc (ng/µl) | VolEl (µL) |
|---|---|---|---|---|---|---|---|---|---|
| ACO 27 | 12 /48 | O3 | 21.09.2023 | 10:07-11:10 | 2-3mm | RNeasy Plus Micro | 10.0 | 90.5 | 20 |
| ACI 27 | 12 /48 | I3 | 21.09.2023 | 10:07-11:10 | 2-3mm | RNeasy Plus Micro | 10.0 | 67.4 | 20 |
| ACO 24 | 12 /48 | O3 | 11.09.2023 | 10:31-11:38 | 2-3mm | RNeasy Plus Micro | 10.0 | 108 | 14 |
| ACI 24 | 12 /48 | I3 | 11.09.2023 | 10:31-11:38 | 2-3mm | RNeasy Plus Micro | 10.0 | 41.3 | 14 |
| ACO 25 | 12 /48 | O3 | 12.09.2023 | 10:00-10:51 | 2-3mm | RNeasy Plus Micro | 10.0 | 62.3 | 20 |
| ACI 25 | 12 /48 | I3 | 12.09.2023 | 10:00-10:51 | 2-3mm | RNeasy Plus Micro | 10.0 | 95.1 | 20 |
| ACO 28 | 8 /32 | O5 | 21.09.2023 | 11:11-11:43 | 5-5.5mm | RNeasy Plus Micro | 9.7 | 263 | 30 |
| ACI 28 | 8 /32 | I5 | 21.09.2023 | 11:11-11:43 | 5-5.5mm | RNeasy Plus Micro | 9.8 | 273 | 30 |
| ACO 23 | 12 /48 | O5 | 07.09.2023 | 10:01-11:07 | 5-5.5mm | RNeasy Plus Micro | 6.8* | 368 | 14 |
| ACI 23 | 12 /48 | I5 | 07.09.2023 | 10:01-11:07 | 5-5.5mm | RNeasy Plus Micro | 6.8* | 387 | 14 |
| ACO 26 | 10 /40 | O5 | 12.09.2023 | 10:55-11:42 | 5-5.5mm | RNeasy Plus Micro | 9.9 | 305 | 30 |
| ACI 26 | 10 /40 | I5 | 12.09.2023 | 10:55-11:42 | 5-5.5mm | RNeasy Plus Micro | 9.8 | 267 | 30 |
| ACO 15 | 3 /12 | O12 | 26.07.2023 | 10:58-11:14 | 11-12mm | RNeasy Plant mini | 9.8 | 51.6 | 30 |
| ACI 15 | 3 /12 | I12 | 26.07.2023 | 10:58-11:14 | 11-12mm | RNeasy Plant mini | 8.8 | 40.2 | 30 |
| ACO 17 | 3 /12 | O12 | 31.07.2023 | 11:23-11:42 | 11-12mm | RNeasy Plant mini | 9.5 | 42.7 | 30 |
| ACI 17 | 3 /12 | I12 | 31.07.2023 | 11:23-11:42 | 11-12mm | RNeasy Plant mini | 9.5 | 45 | 30 |
| ACO 21 | 3 /12 | O12 | 30.08.2023 | 10:25-10:39 | 11-12mm | RNeasy Plant mini | 9.9 | 69.6 | 30 |
| ACI 21 | 3 /12 | I12 | 30.08.2023 | 10:25-10:39 | 11-12mm | RNeasy Plant mini | 9.7 | 39.8 | 30 |
| ACO 13 | 3 /12 | O18 | 25.07.2023 | 10:48-10:58 | 17-18mm | RNeasy Plant mini | 9.3 | 221 | 30 |
| ACI 13 | 3 /12 | I18 | 25.07.2023 | 10:48-10:58 | 17-18mm | RNeasy Plant mini | 9.1 | 169 | 30 |
| ACO 14 | 3 /12 | O18 | 26.07.2023 | 10:37-10:52 | 17-18mm | RNeasy Plant mini | 8.7 | 175 | 30 |
| ACI 14 | 3 /12 | I18 | 26.07.2023 | 10:37-10:52 | 17-18mm | RNeasy Plant mini | 9.7 | 123 | 30 |
| ACO 16 | 3 /12 | O18 | 31.07.2023 | 11:06-11:22 | 17-18mm | RNeasy Plant mini | 10.0 | 316 | 30 |
| ACI 16 | 3 /12 | I18 | 31.07.2023 | 11:06-11:22 | 17-18mm | RNeasy Plant mini | 9.9 | 311 | 30 |

*RINe value lower than true value due to excessive amounts of RNA used during measurement. The values improved to above 9.0 upon dilution
