## Supplementary figures and images for "Insights into the functional and genetic basis of heteranthery in *Arthrostemma ciliatum* Pav. ex D.Don (Melastomataceae)"

### Appendix-2: Developmental series images of flower buds and stamen

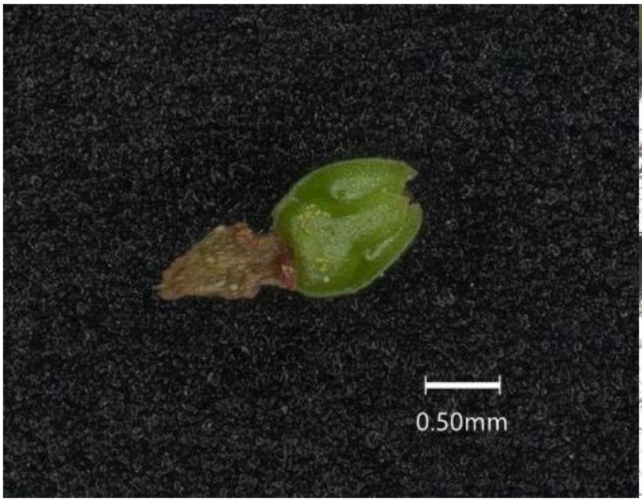

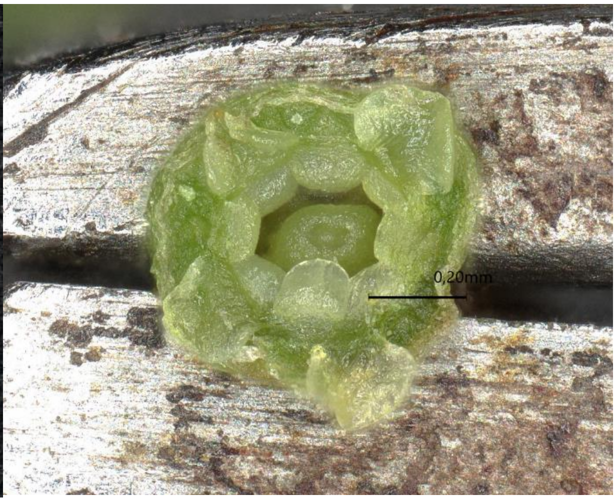

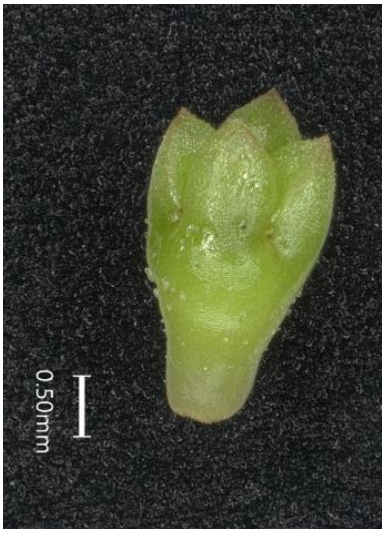

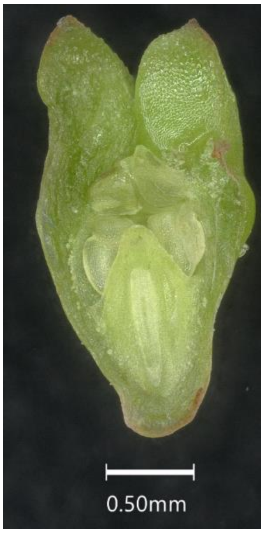

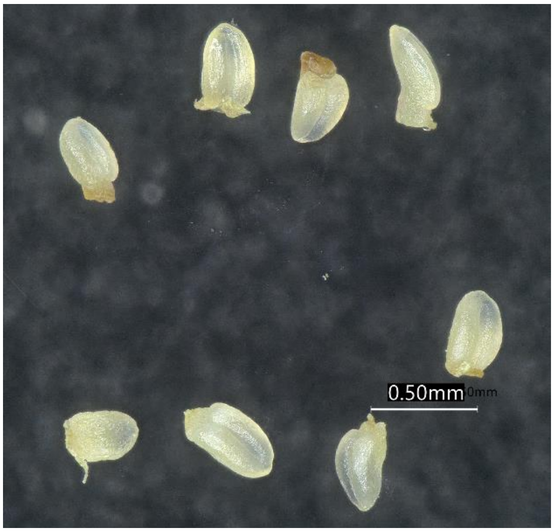

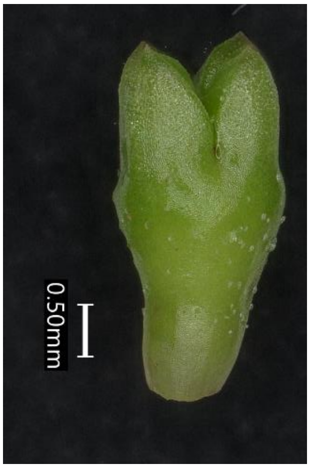

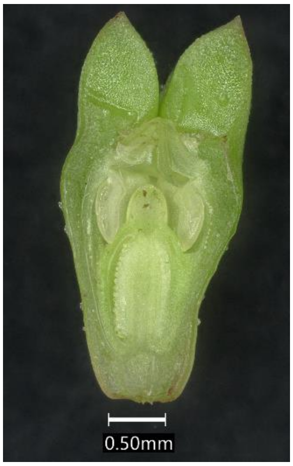

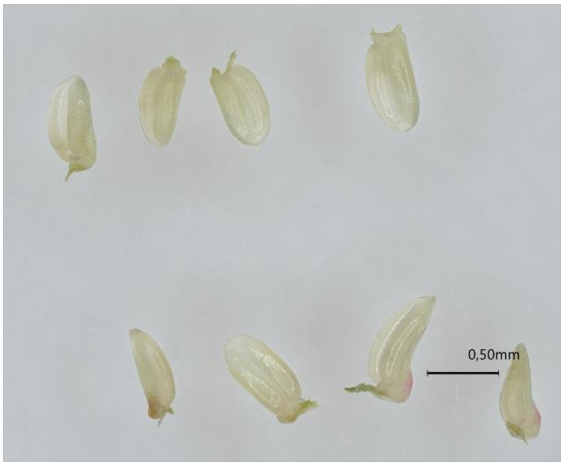

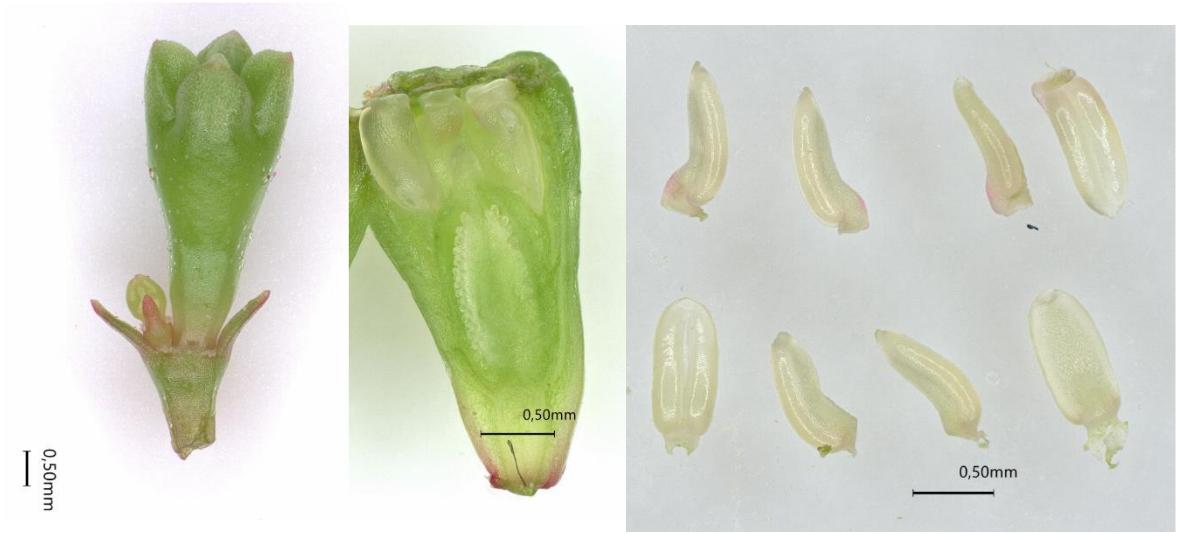

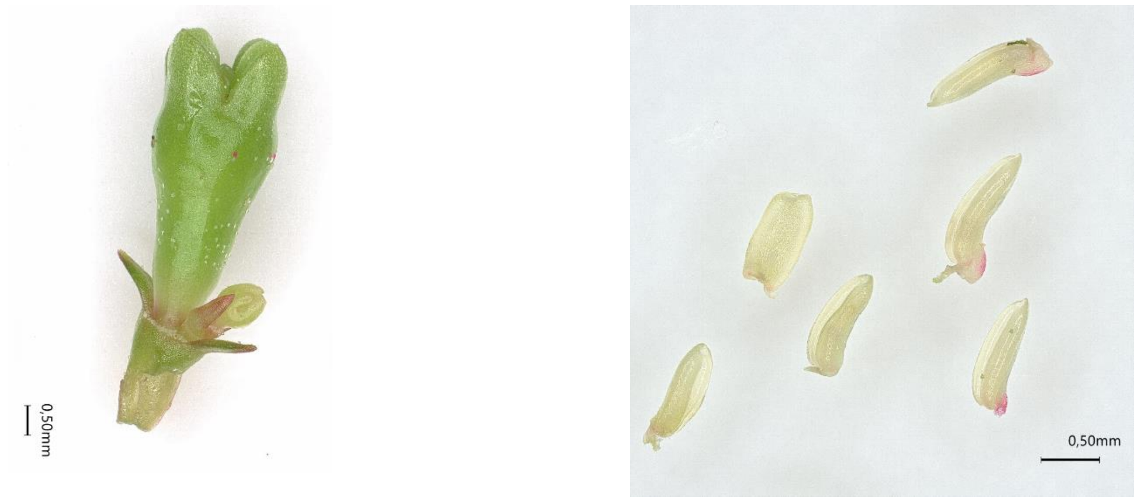

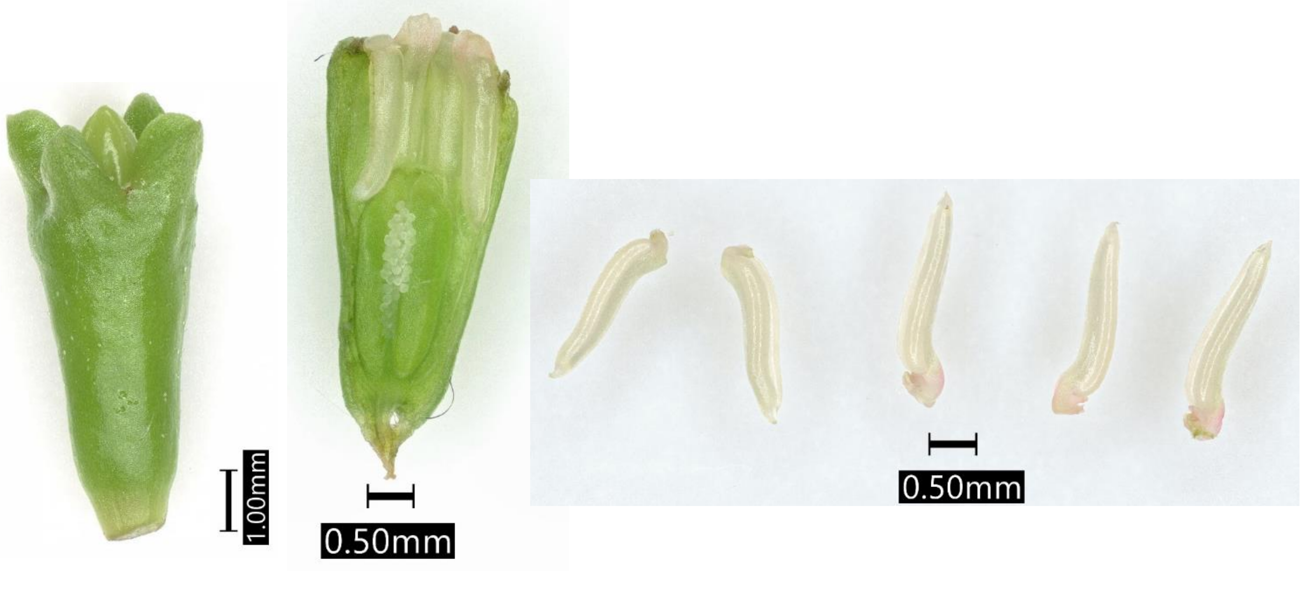

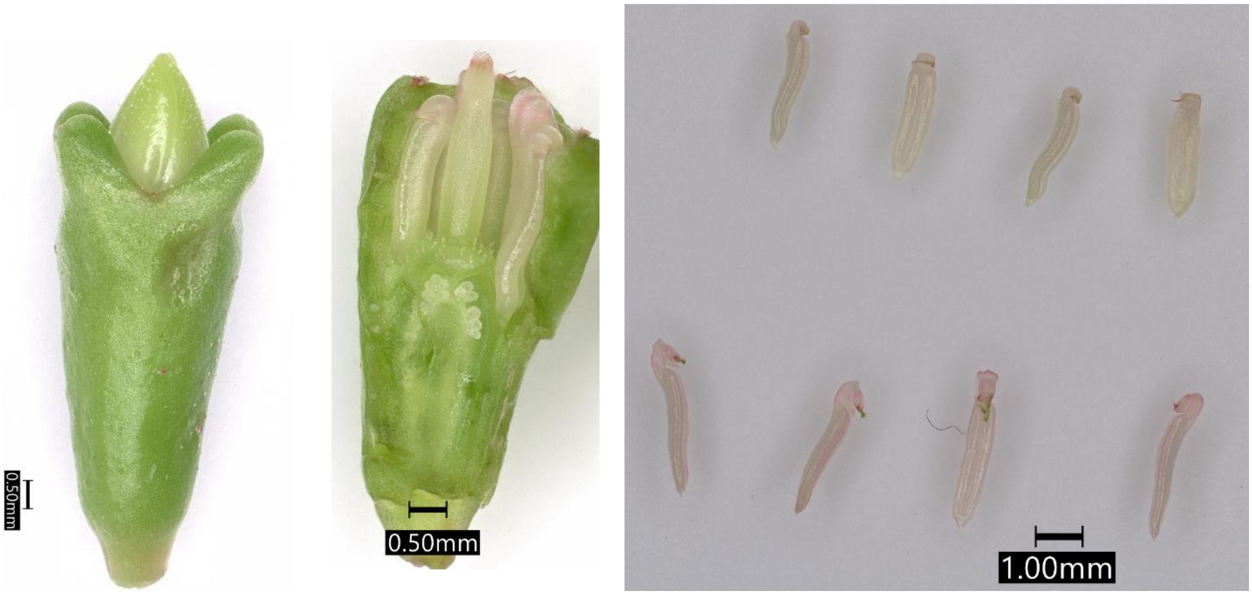

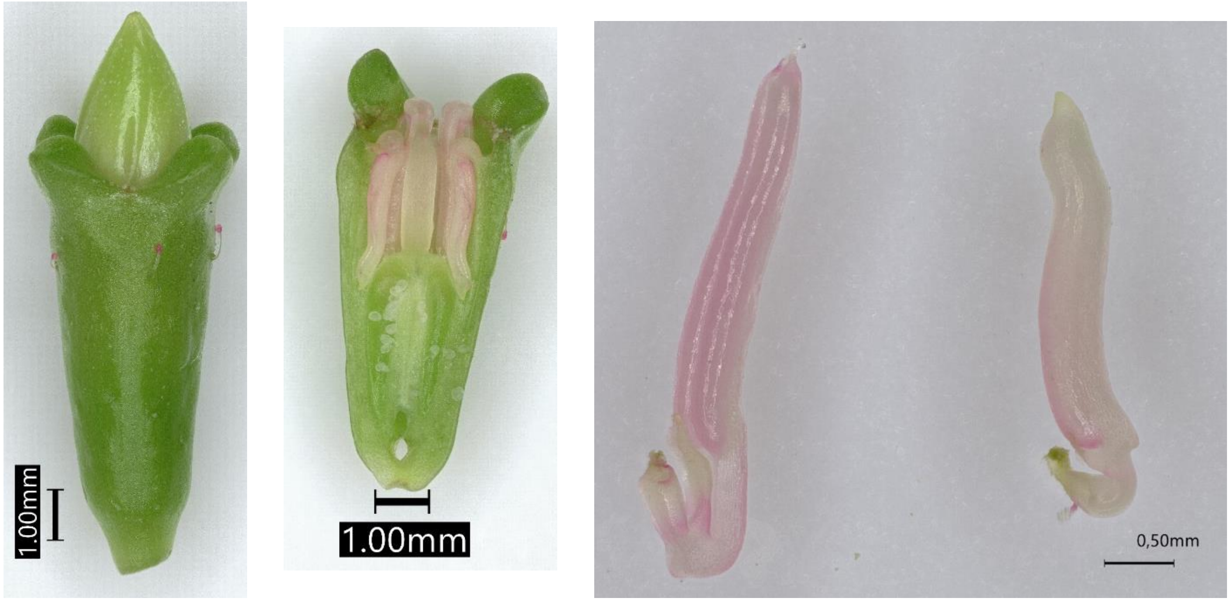

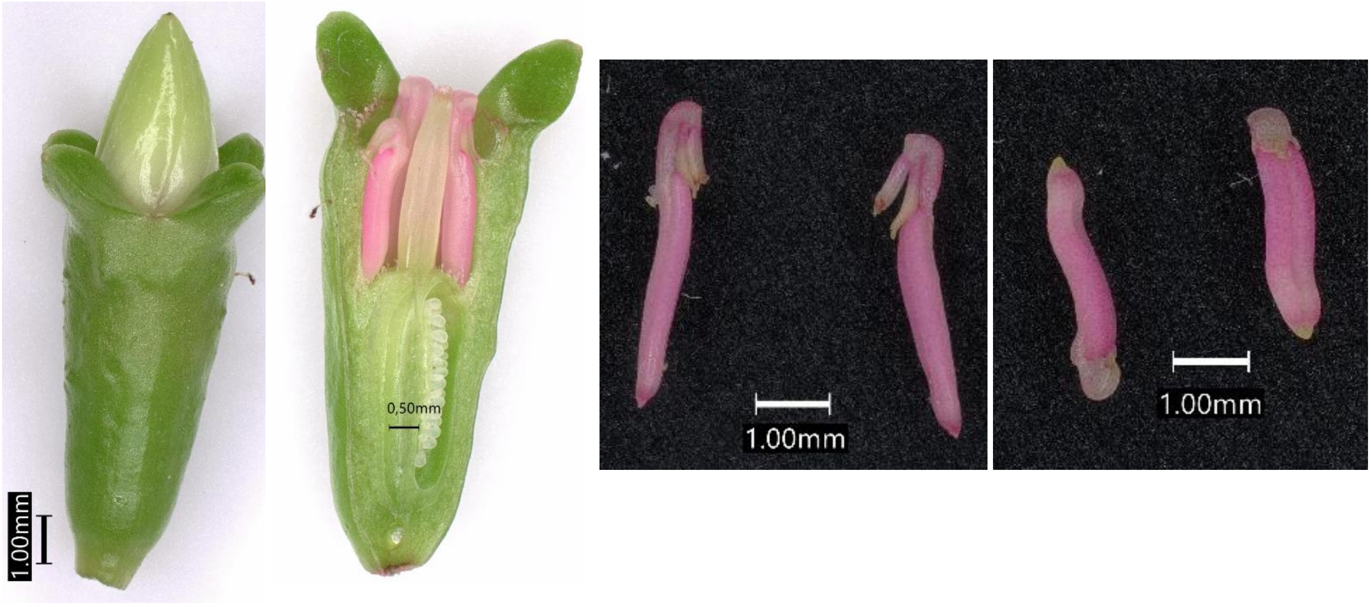

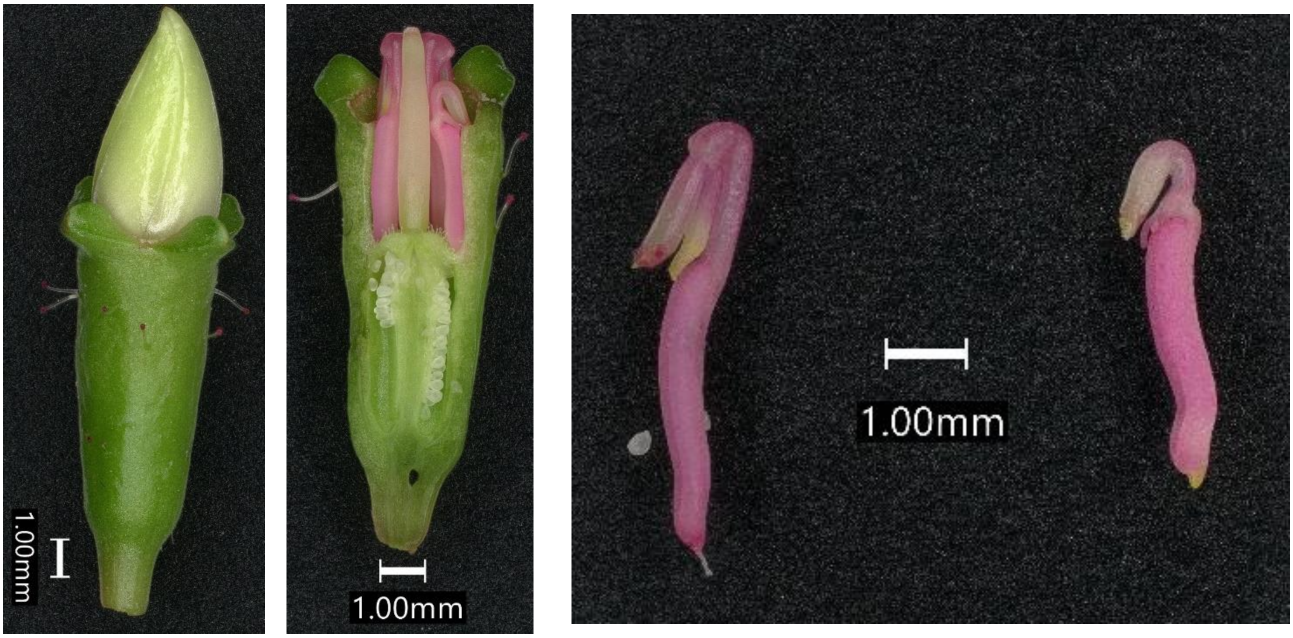

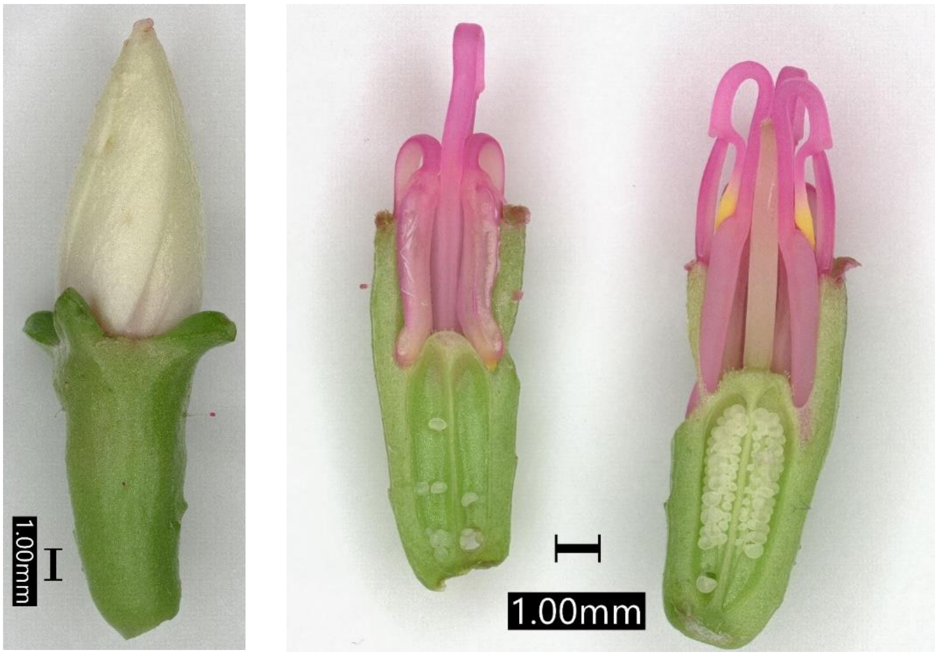

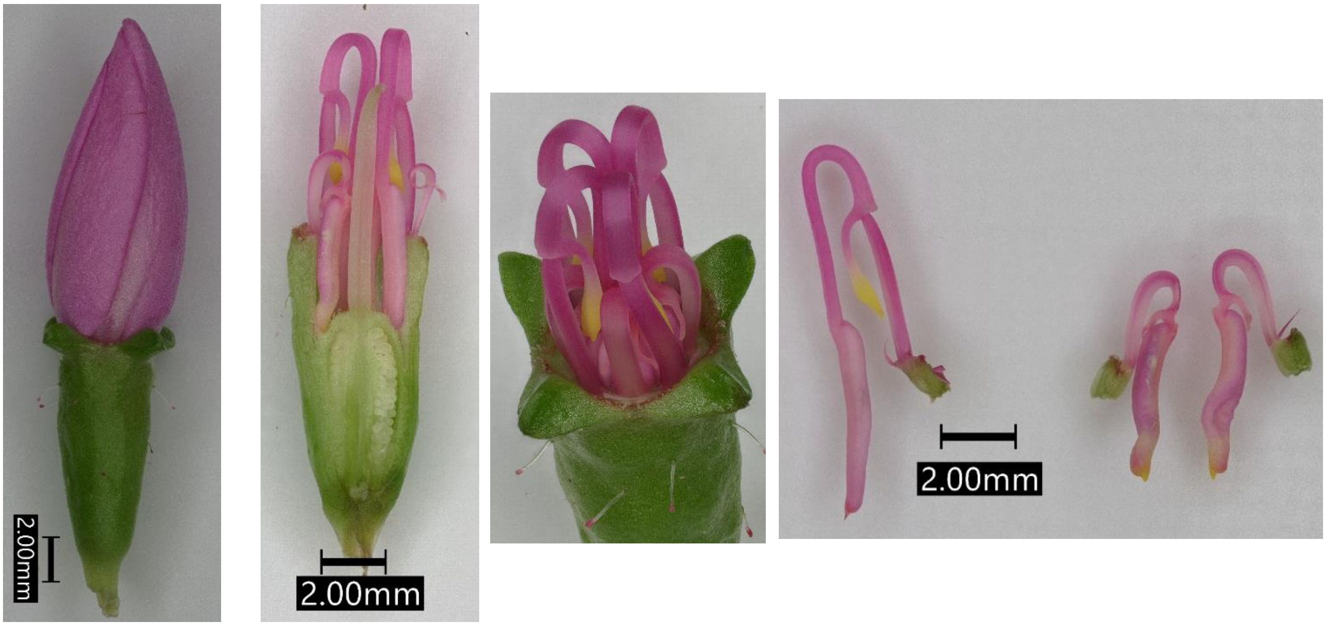

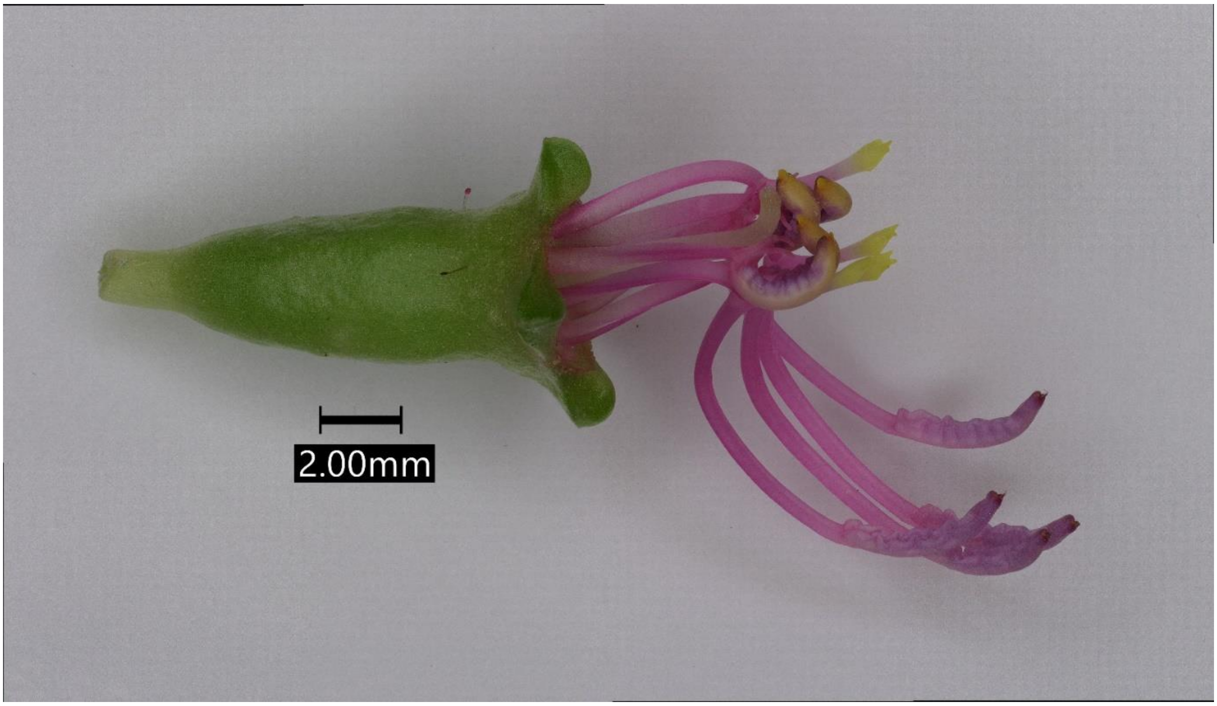

### Appendix-4: Scatterplot of transformed counts from two samples

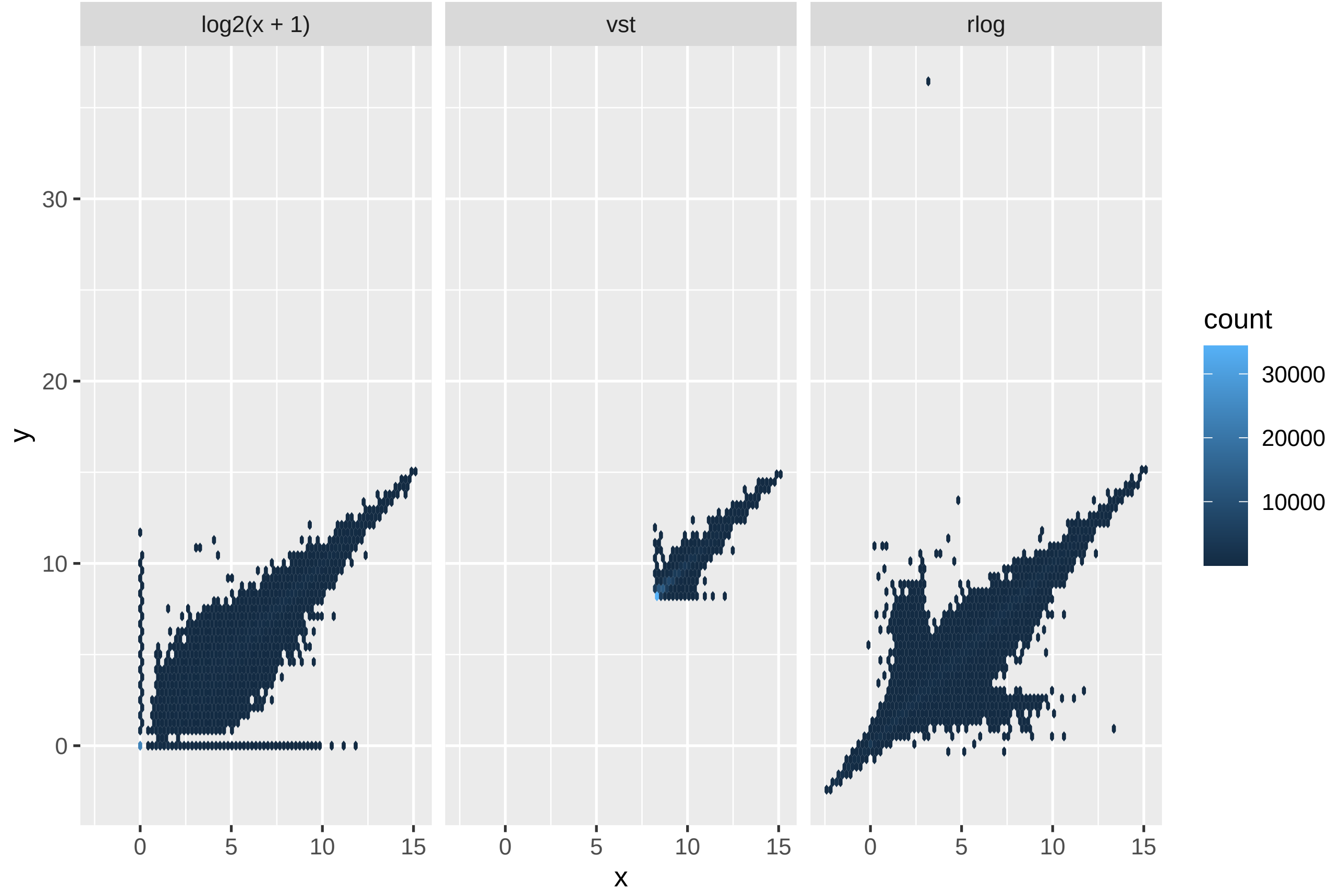

### Appendix-6: Violin plots of theca size by stamen type and flower

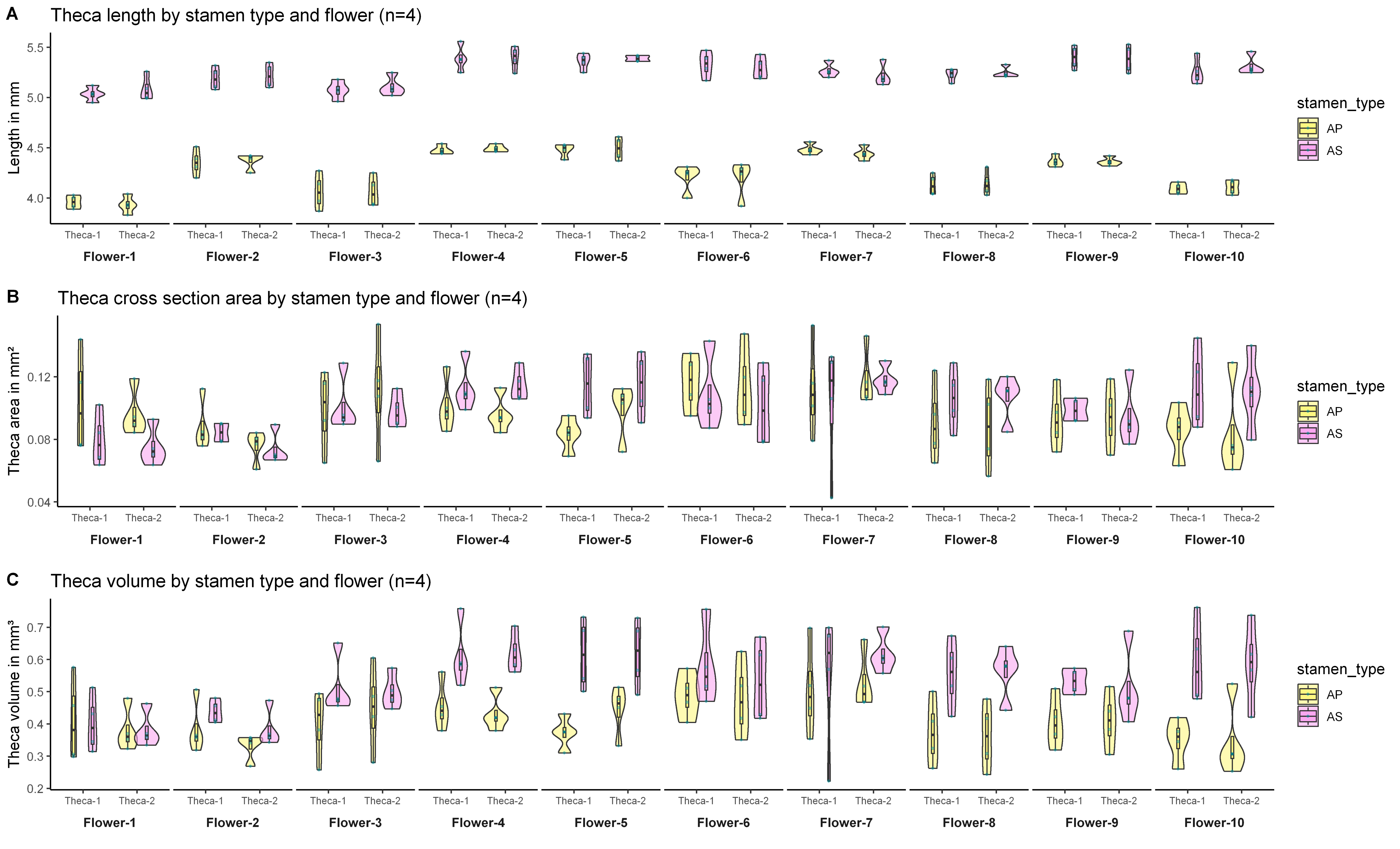

### Appendix-7: Violin plots of length of anthers used for pollen counts and pollen numbers by stamen type

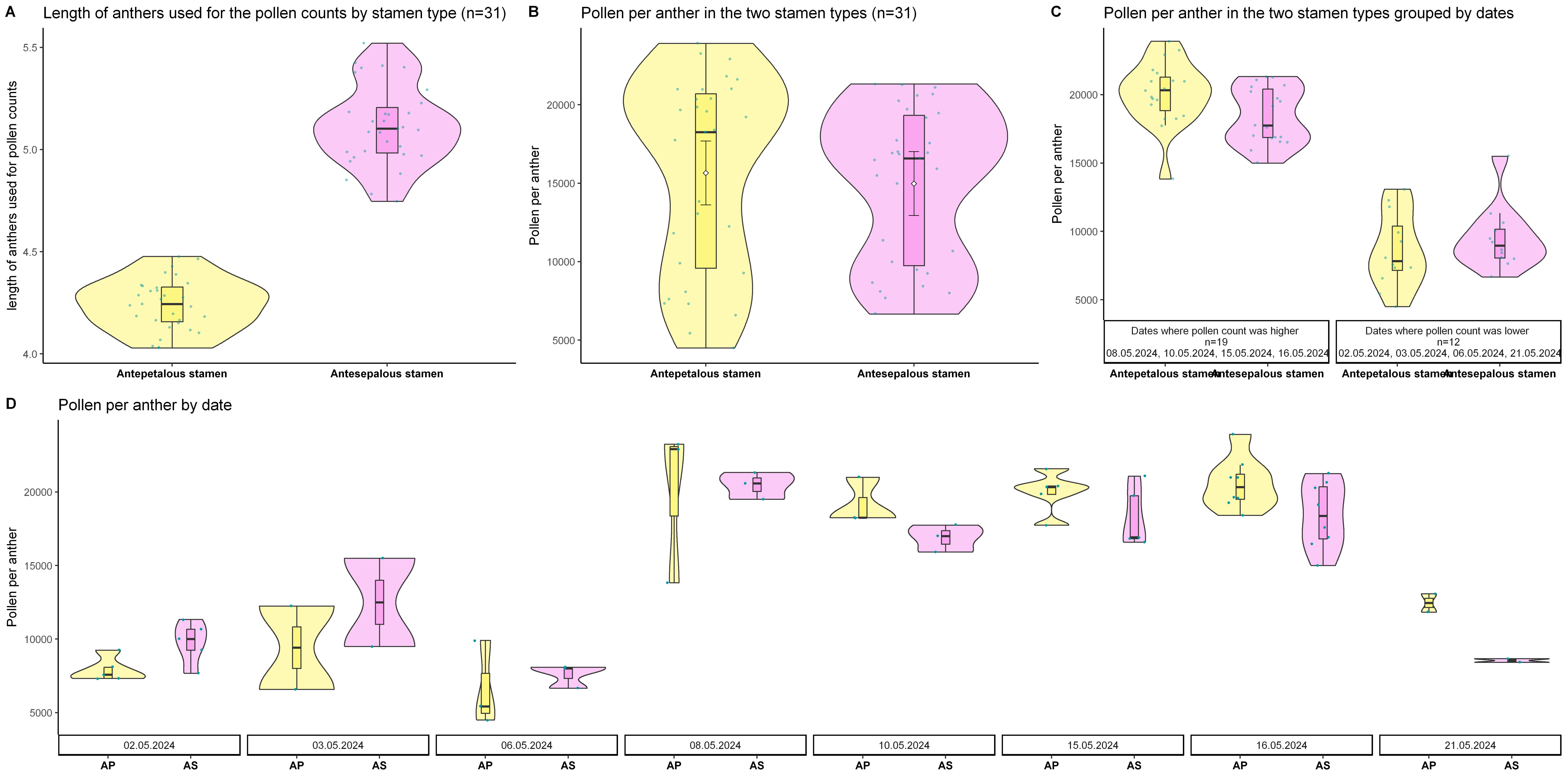

### Appendix-8: Violin plots of pollen size by stamen type and flower

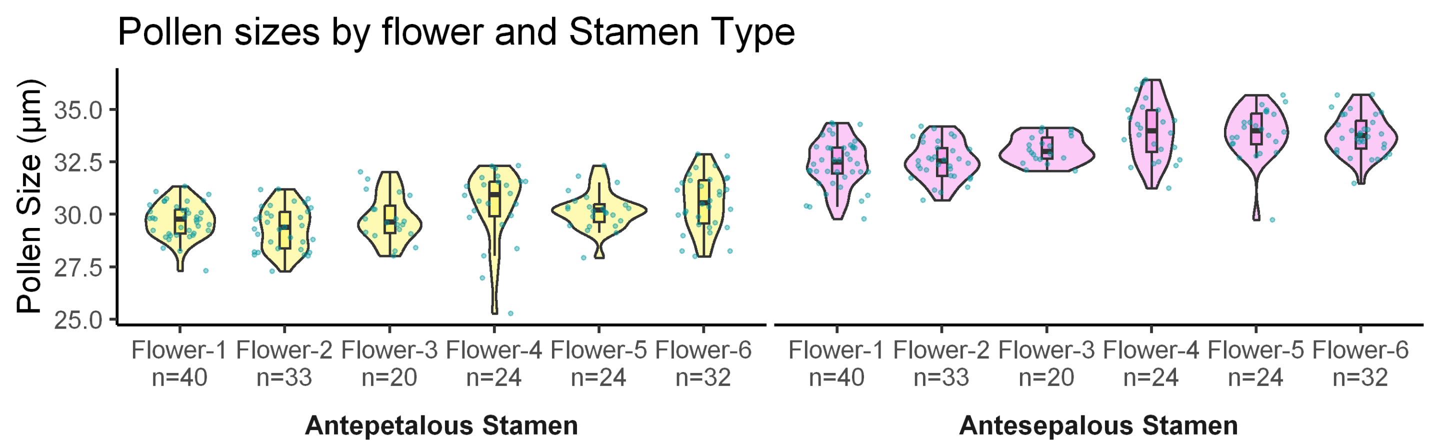

### Appendix-9: Violin plots of pollen germination by stamen types and flower

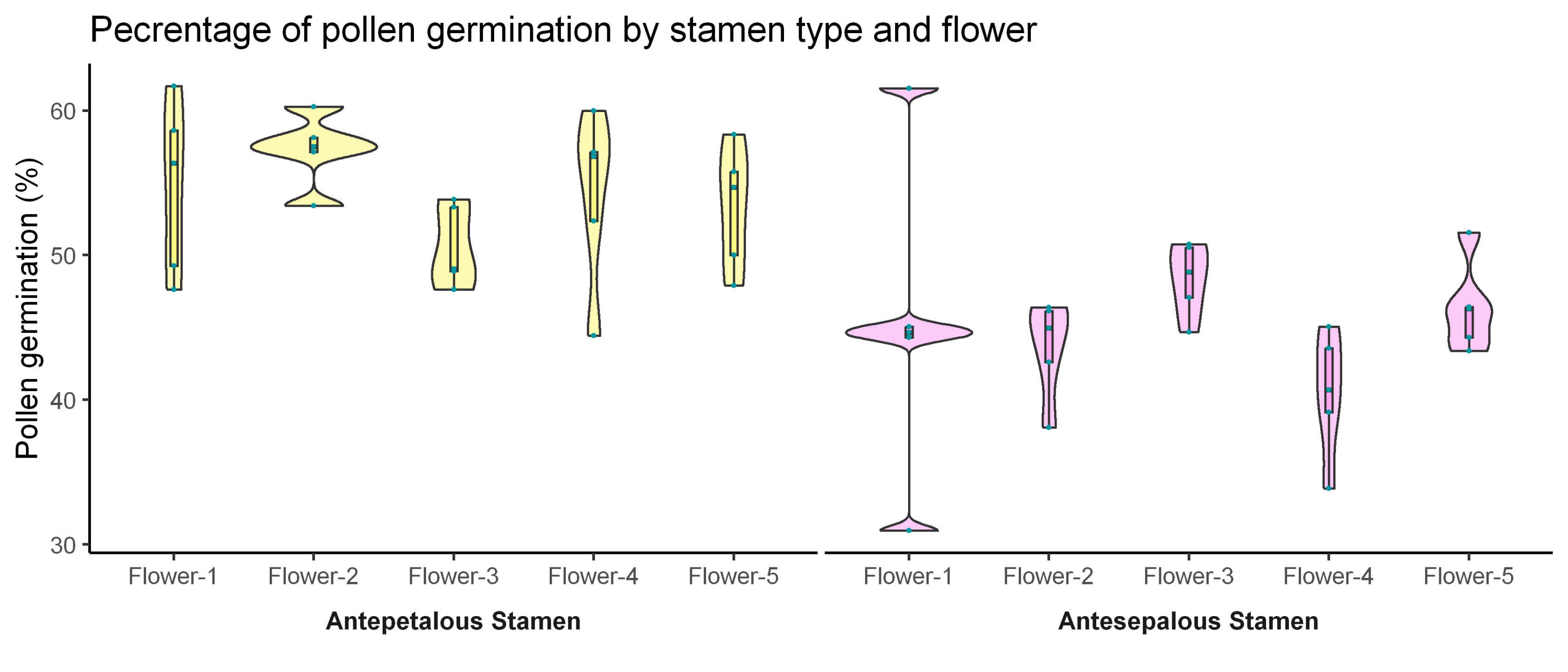
